# Adaptation to marginal habitats at the rear edge provides resilience to future warming

**DOI:** 10.64898/2026.01.30.702836

**Authors:** Antoine Perrier, Laura F. Galloway

## Abstract

Rear-edge populations are typically relicts of glacial refugia that persist in marginal habitats at the warmer edge of species distributions. Persistence likely required adapting to postglacial climate warming. The resulting climate adaptation may affect success under future climate change if adaptative strategies align with future conditions. We explored how rear-edge populations persist in their marginal climates, and whether any adaptations affect responses to expected future warming in the North American herb *Campanula americana*. We raised plants from 23 rear-edge and central populations in three common gardens spanning their natural climatic gradient. We evaluated performance across the life cycle in each garden to identify key life stages underlying climate adaptation at the rear-edge. We then predicted population performance at the key stage under future climates. Local climate adaptation at the rear edge was mainly driven by whether plants were able to initiate bolting, thereby transitioning from vegetative growth to reproduction. Rear-edge populations exhibited high bolting frequency under local warm winters, while more northern populations failed to bolt in these conditions. Bolting is cued by winter cold (vernalization) in the species, but rear-edge climates provide only a few days sufficiently cold for effective vernalization. We found that rear-edge populations have adapted to these conditions by evolving reduced vernalization requirement and sensitivity. Under projected warmer winters, bolting is expected to decline at mid-latitudes, whereas rear-edge populations are predicted to maintain high bolting. Our findings highlight that adaptations that likely allowed rear-edge populations to persist under past warming may also buffer these populations against future warming.

## Introduction

Contemporary climate change is expected to disrupt global species distributions (Lenoir & Svenning, 2015; Rubenstein *et al*., 2023; Lenoir *et al*., 2024; Urban, 2024). In temperate regions, ecological models typically predict that species will shift their ranges towards higher latitudes and elevations to track suitable habitats (Parmesan & Yohe, 2003). High risk of extinction is expected at warmer range limits as future climates may exceed species’ thermal niche limits (Cahill *et al*., 2014). While such responses are already broadly observed under ongoing climate change (e.g. Anderson *et al*., 2025), lags in expected warm-edge contractions are increasingly documented, particularly in plant species (Alexander *et al*., 2018; Vilà-Cabrera *et al*., 2019; Geppert *et al*., 2020; Duchenne *et al*., 2021; Rubenstein *et al*., 2023). Whether warm-edge populations persist or decline under climate change is likely to depend on interactions between ecological factors, e.g. the pace and type of environmental change, and evolutionary factors, e.g. genetic drift and adaptation (Nadeau & Urban, 2019; Aguirre-Liguori *et al*., 2021). Similar interactions shaped warm range limits under past climate change (Hewitt, 2000, 2004; Davis & Shaw, 2001; de Lafontaine *et al*., 2018), and may have lasting evolutionary legacies, influencing contemporary vulnerability and resilience to future warming (Perrier *et al*., 2026b). Studying the outcomes of these past processes, and their consequences under future warming, may help illuminate the observed heterogeneity of warm-edge responses to climate change.

Rear-edge populations, typically glacial relicts persisting at warmer range limits, represent ideal models to study responses to warming (Hampe & Petit, 2005; Perrier *et al*., 2026b). During the Last Glacial Maximum (LGM, ∼20kya; Hughes *et al*., 2013), many temperate species took refuge in unglaciated areas at low latitudes and elevations, then expanded out of these refugia as the earth warmed (i.e., “expanded range”; Hewitt, 2000, 2004). In some species, historic refugial populations persisted more-or-less in place to the present day, forming the “rear edge” of contemporary species distributions (“stable edge” sensu Hampe & Petit, 2005). These populations often occur in habitats that are unsuitable relative to the species niche (Vilà-Cabrera *et al*., 2019; Bontrager *et al.,* 2021), suggesting they persisted by gradually adapting to the selection imposed by long-term warming (Bontrager *et al*., 2021; Perrier *et al*., 2026b; a). Rear-edge populations may thus serve as a natural laboratory to study adaptations that were successful under past warming, and the effect of such adaptations on responses to projected future warming.

Adaptations that are likely to have evolved in response to past warming can be inferred by studying how rear-edge populations have adapted to their local climate (hereafter “local adaptation”). In particular, investigating performance at key life stages (e.g. survival, life-stage transitions, reproduction), and characterizing how performance at these stages varies with environmental conditions, can help identify the adaptations (Garnier *et al*., 2015) underlying persistence at the rear edge. The relationship between environment and performance at key life stages can also be used to predict population- and species-level responses to future climate change (e.g. Pacifici *et al*., 2017; Jackson *et al*., 2022). Rear-edge populations have been shown to harbor distinct trait variation associated with warmer climates, including morphological and physiological traits linked to abiotic stress tolerance (Mathiasen & Premoli, 2016; Saada *et al*., 2016; King *et al*., 2019; Ghouil *et al*., 2020), and environmentally cued traits regulating life history (Pelletier & de Lafontaine, 2023; Perrier *et al*., 2025). However, empirical studies of local adaptation at the rear edge are scarce (Perrier *et al*., 2026b), and the life stages key for persistence in novel rear-edge climates, and the types of adaptation that result in high performance, remain largely untested.

Adaptation to past warming may directly affect population fitness under future warming, thereby influencing whether warm edges contract or remain stable. Rear-edge populations are thought to be at high risk of extinction under predicted future warming as they often occur at a species’ warmer ecological or physiological limit (Cahill *et al*., 2014; Nadeau & Urban, 2019; but see Vilà-Cabrera *et al*., 2019). Some studies support this expectation finding rear-edge populations more vulnerable to future warming than other parts of the range (Sánchez-Salguero *et al*., 2017; Assis *et al*., 2018; Anderson *et al*., 2025). However, other studies find that rear-edge populations may be more resilient (reviewed in Vilà-Cabrera *et al*., 2019; e.g. DeMarche *et al*., 2018; Vilà-Cabrera & Jump, 2019). This resilience may be linked to adaptations that enable persistence in the warm rear-edge climate, for example greater resistance/tolerance of environmental stress (Nicotra *et al*., 2010; Franks *et al*., 2014; Hamann *et al*., 2018; Kristensen *et al*., 2020), escape from stress (Kooyers, 2015), or fitness that is decoupled from environmental change (Anderson, 2016). However, how potential response to future warming relates to adaptations expected to have enabled persistence under past warming has rarely been tested.

In this study, we evaluate whether adaptations that likely allowed rear-edge populations to persist through past warming improve their expected response to future warming. We address this in the North American plant *Campanula americana,* where a previous multisite common-garden experiment revealed strong local adaptation at the rear edge (Perrier *et al*., 2026a). Building upon that result, we investigated how rear-edge populations are adapted to their local warmer climates (plants and data as in Perrier *et al*., 2026a). Specifically, we ask: in rear-edge climates (Q1) which life stages are key to achieving high fitness, and (Q2) what adaptation allows high performance at these life stages? After identifying a key adaptation that allows persistence at the rear edge, we use forecasts of future climate scenarios to ask: (Q3) Do adaptations that are expected to have allowed rear-edge populations to persist under past warming improve their predicted performance under future climates? We focus on southern rear-edge populations of the species, as these show the strongest local adaptation (Perrier *et al*., 2026a), but also include populations from the rest of the range for comparison. Greater performance of rear-edge populations under projected warming relative to the rest of the range would support resilience conferred by adaptation to past warming. Overall, our study links adaptations expected to have evolved in response to past climate change to resilience under future climate change, providing insight into how such evolution may mediate range contraction or stability of warmer range limits under ongoing climate change.

## Material and Methods

### Study system

*Campanula americana* is an outcrossing monocarpic herb found in open forests across the eastern USA (Fig. 1A). Seeds are produced in the fall, with many germinating shortly after dispersal. Seedlings overwinter as vegetative rosettes (Baskin & Baskin, 1984; Galloway, 2002). In *C. americana*, the transition to reproduction, i.e. bolting, is cued once a critical duration of cold exposure is reached (*i.e.* vernalization; Perrier *et al*., 2025). Bolting is initiated in spring with the elongation of rosettes. After several months of growth, flowering begins mid-summer and lasts ca. five weeks with fruits taking a month to mature.

**Figure 1:**
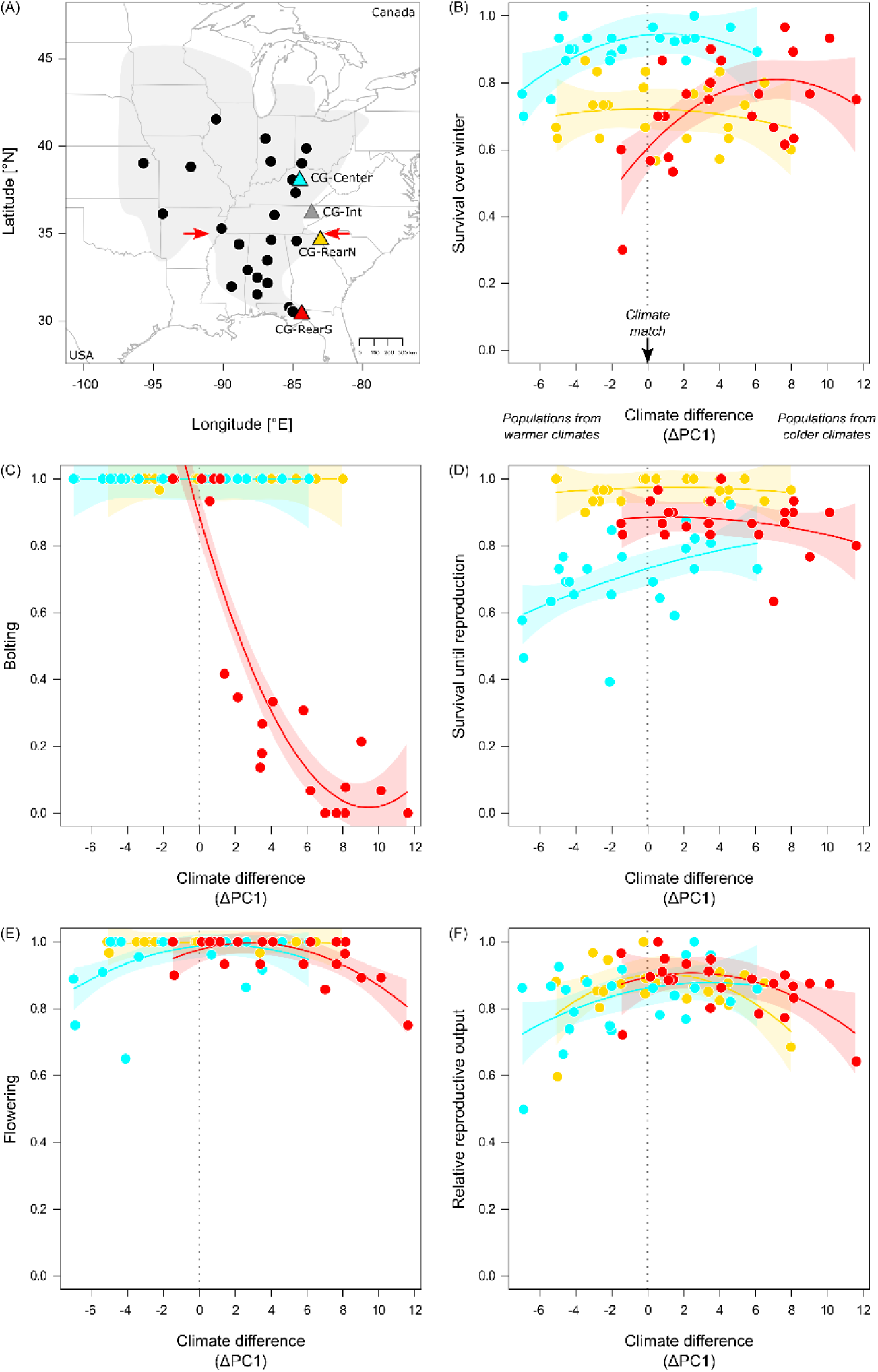
Bolting is the key trait underlying local adaptation at the rear edge. **(A)** Sampling of *C. americana* populations (dots) and common gardens (triangles), with the range of the Western lineage shaded (light gray). Red arrows indicate the latitudinal delimitation of the rear edge. CG- Int (gray) was not included in the analysis of local adaptation due to premature death. **(B-F)** Relative performance based on life stages of populations (dots) raised in each common garden (distinguished by color as in A). Solid lines represent the relationship between relative trait performance and the climate difference between each garden and population’s location of origin (ΔPC1), estimated for each garden, with the 95% CI indicated in shading (test statistics: Table 1, S12). The dashed line (climate difference = 0) indicates a climate match between the population’s origin and the garden.

**Table 1:** Local adaptation across the life cycle.

| Life stage | Common garden<br>(CG) | Climate difference<br>($\Delta PC1 + \Delta PC1^2$) | CG *<br>($\Delta PC1 + \Delta PC1^2$) | $R^2_m$ $R^2_c$ |
| --- | --- | --- | --- | --- |
| | $X^2$ | $X^2$ | $X^2$ | |
| Survival over winter | **56.84** *** | **9.46** ** | **17.31** *** | 0.52 0.68 |
| Bolting | **12.19** ** | **91.73** *** | **140.59** *** | 0.92 0.93 |
| Survival until reproduction | **74.54** *** | 1.33 | 5.06 | 0.67 0.76 |
| Flowering | **8.23** * | 5.68 | 4.97 | 0.34 0.50 |
| Reproductive output | 0.74 | 3.61 | 6.22 | 0.25 0.75 |

Populations belong to three geographically distinct genetic clades (Barnard-Kubow *et al*., 2015; Lamb *et al*., 2024). We focus on the larger “Western” clade (∼ 80% of all populations) found at low elevation west of the Appalachian Mountains. The rear edge of the Western clade is geographically defined as the lower latitudinal third of the range (< 35° N), east of the Mississippi (Fig. 1A, Perrier *et al*., 2026c). Populations in this area overlap putative glacial refugia, are ancestral to the rest of the clade (Barnard-Kubow *et al*., 2015; Perrier *et al*., 2026c), and are highly differentiated from each other over short geographic distances (Perrier *et al*., 2026c), and thus likely represent descendants of refugial populations that persisted more or less in place. The history of postglacial warming varies across the rear edge, with the southernmost populations (∼31° N, hereafter “southern rear edge”) having undergone stronger changes in climate than populations at the northern edge of the former refugia (∼35° N, hereafter “northern rear edge”; Perrier *et al*., 2026c). The rest of the clade’s range resulted from north- and westward postglacial range expansion (Koski *et al*., 2019; Prior *et al*., 2020), with populations likely tracking shifts in suitable habitats and experiencing little warming (Perrier *et al*., 2026c).

*Campanula americana* spans a large climatic range, from subtropical areas near the Gulf coast to the colder Great Lakes region. Contemporary rear-edge habitats occur in subtropical climates substantially warmer than the continental climates elsewhere in the range (Perrier *et al*., 2025), and with low suitability relative to the species niche (Barnard-Kubow *et al*., 2015, Perrier *et al.,* 2026c). However, southern rear-edge populations have adapted to thrive in these warmer conditions, showing stronger local adaptation than populations elsewhere in the range (Perrier *et al*., 2026a).

The combination of a history of sustained warming with strong local adaptation suggests southern rear-edge populations persisted in place by adapting to past warming. This makes them ideal for exploring whether adaptations that likely evolved in response to past warming may affect responses to future warming. Our focus is therefore these populations. In these populations, climatic differences compared to the rest of the range, particularly warmer winters, have been associated with reduced vernalization requirements and phenological shifts (Perrier *et al*., 2025). However, the life stage key to persistence in these warm climates, the underlying adaptions, and the potential of these adaptations for mediating the southern rear-edge populations’ response to future warming, remain unknown.

### Q1: Which life stages are key for climate adaptation at the southern rear edge?

In a previous common garden experiment, we determined that southern rear-edge populations showed strong local adaptation for lifetime fitness (Perrier *et al*., 2026a; also variation in flowering phenology, Perrier *et al*., 2025), but the source of this adaptation was not characterized. Using the data from that experiment, we identify the life stage underlying the climate-specific adaptation that allows southern rear-edge populations to thrive in habitats that are unsuitably warm compared to the rest of the range (Perrier *et al*., 2026c).

#### Experimental design

We selected 23 source populations (Fig. 1A, Table S1) across a latitudinal gradient from the southernmost edge to the range center (30.6° N to 41.6° N). This sampling allowed us to study adaptation in the southern rear-edge populations by comparing them with populations that have experienced less warming and occupy habitats more typical of the species. For each source population, we collected seeds from an average of 27 maternal plants (hereafter seed families) in the field in late summer 2020 and 2021. We propagated them for one generation in common greenhouse conditions (details in Perrier *et al*., 2026a). We used the greenhouse-produced seeds for 20 populations, and field-collected seeds for four populations that lacked greenhouse seeds in the common garden experiment (Table S1). We raised plants from each source population in each of three common gardens located along an 850 km latitudinal gradient (CG, Fig. 1A, Table S2A). The gardens represent conditions in the southern rear edge (CG-RearS, Tallahassee FL, 30.5°N), the northern rear edge near the northern extent of the former refugia (CG-RearN, Clemson SC, 34.7°N), and the center of the range with a climate more typical of the species (CG-Center, Lexington KY, 38.1°N). CG-RearN and CG-Center serve as comparisons to identify adaptations that are specific to southern rear-edge conditions. A fourth garden, intermediate between the northern rear edge and center gardens (CG-Int, Blaine TN, 36.2°N), was only included in Q2 and Q3 (below) due to premature death of most individuals during the reproductive period.

Plants were raised in two successive cohorts that together capture the species’ life cycle (Fig. S1, see Perrier *et al*., 2026a). In the “winter cohort”, we study the effect of winter conditions on seedling survival and cuing of reproduction (bolting). Winter-cohort seeds were germinated in controlled conditions simulating fall. Seedlings were then transplanted to each garden in fall 2023, before the onset of winter (Table S2B). Plants were tracked through bolting (late spring 2024). In the “summer cohort”, we study how spring and summer conditions affect survival and reproduction of seedlings that were adequately cued for reproduction. Summer cohort seedlings were germinated in controlled conditions simulating fall (2022), vernalized in a simulated winter to cue reproduction, then transplanted at the end of natural winter into each garden (spring 2023; Table S2A). Plants grew until late summer and were harvested when most had mature fruit. We use two cohorts because if many fall seedlings did not survive the winter or failed to transition to reproduction, replication would be too low to capture variation in reproductive traits in summer.

Within each of the three gardens, we raised 30 individuals per source population for each cohort (15 seed families x 2 replicates per family), for a total of 4140 individuals (30 individuals/population x 23 populations x 2 cohorts x 3 gardens). In each planting site, the garden was placed in an area similar to the natural habitat of the species. We aimed to limit biotic interactions on plant performance to improve detection of local adaptation linked to climate (Hargreaves *et al*., 2020). We therefore fenced gardens to prevent browsing by large herbivores. We also removed aboveground portions of resident plants at the time of transplant to facilitate establishment of seedlings and reduce competition. No further intervention was performed. We placed one data logger (iButton®; Maxim Integrated Products Inc., San Jose, CA, USA) in the shade in each garden, 1.5 m aboveground, to monitor air temperature hourly for the duration of the experiment.

#### Performance across the life cycle

We recorded plant performance at critical vegetative and reproductive stages of the life cycle (data as in Perrier *et al*., 2026a). We visited the winter cohort two times when plants were bolting, in early spring (March) and late spring (May-June, Table S2), and the summer cohort two or three times when plants were reproducing (July and September). At each visit, we assessed survival and transition to bolting for each individual. For the summer cohort, we also recorded transition to flowering (production of at least one bud) and at the final visit collected plants to record reproductive output, assessed as the total number of fruits, flowers and buds.

#### Climate difference between common gardens and populations

We determined relevant climate data for source populations and common gardens. We first generated 14 variables that captured climate variation (mean, minimum and maximum temperature, precipitation, number of days of frost and vernalization) at specific life stages, and five bioclimate variables that reflected annual temperature and precipitation seasonality (Table S3A, B). For source populations, we used data from the PRISM database (2018–2022, https://prism.oregonstate.edu, accessed 13/05/2024) to represent typical conditions experienced by populations. For gardens, we used data recorded during the experiment by dataloggers and nearby weather stations to reflect conditions experienced by experimental plants. While the experiment captured climate only in one year, those garden climates were generally comparable to average conditions in nearby populations, with the center garden being slightly warmer than expected for its latitude (Table S3B, Fig. S2).

For each source population in each common garden, we quantified the difference between climate conditions experienced in the garden and at the population’s location of origin (following Perrier *et al*., 2026a). Preliminary analyses (not shown) revealed that many climate variables were correlated across *C. americana*’s range, especially temperature variables. We therefore performed a PCA on the population climate variables to reduce dimensionality and autocorrelation. We found variation in climate across the sampled populations was mainly associated with winter and spring temperature (PC1, 76.65% of variance explained), with PC1 capturing climate differences between the southern rear edge and the rest of the range (Table S3B, Fig. S2). To a lesser extent, precipitation also contributed to variation in climate (PC2, 8.81% of variance explained, Table S3C, Fig. S2), largely capturing variation among central populations. Therefore, we focused on PC1 to characterize climate variation among populations and garden sites (cf. Landoni *et al*., 2024; Anderson *et al*., 2025). We calculated the climate difference between the garden and a population’s home habitat as ΔPC1 = PC1_garden_ – PC1_population location-of-origin_ for each source population in each garden, where 0 indicates climate matching (i.e. the population’s climate is “local” to the garden). This measure of climate difference allowed us to distinguish whether populations experience warmer (ΔPC1 > 0) or colder (ΔPC1 < 0) conditions in the garden than at their location of origin.

#### Which life stage shows the strongest local adaptation in southern rear-edge populations?

We first identified life stages where patterns of population performance were consistent with local adaptation in the southern rear-edge garden and show stronger local adaptation in that garden. We assessed performance at five life stages (Table S4), including: *Survival over winter* (binary), survival of seedlings in the winter cohort from fall transplant until the early spring visit. *Survival until reproduction* (binary), the survival of plants in the summer cohort from spring transplant through reproduction in summer (or the end of the experiment). *Bolting* (binary), the transition to bolting of surviving winter-cohort plants indicating successful reproductive cuing by vernalization, assessed in early and late spring visits. *Flowering* (binary), the transition to flowering of summer- cohort plants. *Reproductive output,* the sum of fruits, flowers and buds of flowering summer-cohort plants. For analysis, we averaged performance at each life stage, first at the seed-family level then at the population level in each garden. Family-level reproductive output was log10 transformed to achieve normality before calculating population averages. Population-level reproductive output was divided by the maximum value within each garden to account for variation in garden quality, and to scale reproductive output between 0 and 1 to match the binary variables. Population-level data was used in analyses as it matches the population-level data for lifetime fitness (see below); preliminary analyses of individual-level data gave similar results (not shown).

For each life stage, we tested whether population performance depended on the climate difference between the population’s origin and each garden. If a life stage contributes to local climate adaptation, we expect performance to be highest in populations where the climate at their location of origin matches the garden’s climate (ΔPC1 = 0), and to decline with increasing climate difference (i.e. following a local vs foreign framework, Kawecki & Ebert, 2004). Stronger declines in performance indicate stronger local adaptation. Variation in population-average performance was tested for each life stage in hierarchical mixed-effect models assuming a normal distribution (following Perrier *et al*., 2026a), using the packages *lme4* (Bates *et al*., 2015) and *blme* (Dorie *et al.,* 2026) in R (R Core Team, 2026). In each model, we included fixed effects of common garden (three levels), climate difference (ΔPC1), and their interaction. Climate difference was modeled with linear and quadratic components (ΔPC1 + ΔPC1^2^). This allowed us to capture cases where performance declines with both positive and negative mismatch between population origin and garden climate, as well as cases where performance patterns were closer to linear in some gardens while quadratic in others. The interaction between garden and ΔPC1 + ΔPC1^2^ was included to test whether the strength or shape of local adaptation varied among gardens. Population was included as a random effect. For each model (and in all subsequent analyses), we tested and confirmed model assumptions.

For life stages where performance varied with climate difference (significant effect of ΔPC1 + ΔPC1^2^), and where this relationship varied among gardens (significant interaction), we characterized patterns of performance within each garden to ensure they are consistent with local adaptation (highest performance close to ΔPC1=0). We then identified which of the life stages conferred the strongest local adaptation in the southern rear-edge garden (steepest decline in performance with climate difference) using post hoc tests (R package *emmeans*, Lenth *et al*., 2025). We also tested variation in performance of each life stage with only climatic variables relevant to that life stage. However, because climatic variation across the range is mostly driven by temperature, and seasonal temperature values across locations are correlated, these analyses yielded similar results to those using PC1. Only analyses with PC1 were retained for simplicity.

#### Which life stage contributes the most to local adaptation in the southern rear edge?

We used the measure of *Lifetime fitness* calculated across the two cohorts in Perrier et al. (2026a). This fitness measure was the product of all five performance estimates for each stage, averaged at the seed-family level in each garden (the lowest replication level common to both cohorts). For a few family–common garden combinations (63 of 1035, ∼6%), life-stage performance data in either cohort was missing (43 summer cohort, 20 winter cohort) because seed families used in the first cohort did not have enough seeds for the second cohort. Missing seed families were replaced with other families from the same population. For each of these family–common garden combinations, we used the average performance of that population in the garden for the life stages with missing performance data. Seed-family level lifetime fitness was log10 transformed (after adding +1 so that all values were positive), averaged at the population level in each garden to reduce zero-inflation, and finally standardized by the highest fitness in each garden to scale fitness between 0 and 1, matching the scale of performance estimates of the individual life stages.

We characterized which life stage contributed the most to the local adaptation observed at the southern rear edge (Perrier et al. 2026a) by identifying the life stage that most influenced lifetime fitness in that garden. To do this, we regressed lifetime fitness in a linear model against performance of all life stages in the southern rear-edge garden. For comparison, we repeated the analysis for the other two gardens. For CG-Center, bolting was excluded as all plants bolted. Correlations in performance among the life stages and the variance inflation factors (VIF) were generally low (Table S5, S6). We then computed the partial *R*^2^ for each life stage using the function *partial_r2* in the R package *sensemakr* (Cinelli & Hazlett, 2020). The life stage that contributed the most to local adaptation in each garden was defined as the one with a partial *R*^2^ higher than 0.5. We complemented this approach with a leave-one-out analysis for each garden. To do this we ran the same model but iteratively removed a single life stage, then calculated the difference between the AICc value of the model without that life stage and the AICc value of the model containing all life stages (ΔAICc). The life stage that led to the highest ΔAICc was deemed the greatest contributor to local adaptation.

### Q2: What adaptations allow high performance in southern rear-edge conditions?

The results from Q1 indicated that whether plants bolted was key to achieve high fitness in southern rear-edge conditions (see Results). In *C. americana*, the probability of bolting is tightly linked to the duration of cold exposure in winter, with reduced vernalization length leading to lower bolting probability (Perrier *et al*., 2025). In warm rear-edge conditions, adaptation likely involves reducing the amount of cold needed to enable bolting (i.e., vernalization requirement) under short winters, and/or altering the sensitivity of bolting probability to variation in winter duration (vernalization sensitivity). We tested these hypotheses by fitting rection norms for bolting probability using the range of winter conditions each population experienced in the common gardens (Q1). We then used these reaction norms to test whether southern rear-edge populations differ in their vernalization requirement and sensitivity relative to populations in the rest of the range. Variables estimated for these analyses are summarized in Table S7A.

#### Bolting probability under different winter lengths

In *C. americana*, bolting only occurs after vernalization requirements have been met (Perrier *et al*., 2025). As described in Q1, we scored bolting in early and late spring in the common garden experiment, resulting in individual-level estimates of *Bolting* (binary) under short and long vernalization lengths for each of the four gardens (eight total, including CG-Int). For the 12 most southern populations with the shortest winters, we grew an additional cohort of seedlings without vernalization to extend the lower end of the range of vernalization length. Seedlings for this “no vernalization” cohort were germinated in standard conditions indoors, and then moved directly to a greenhouse with warm temperatures and long days without the intermediate vernalization step (conditions described in Perrier *et al*. 2026a). Bolting was scored for ca. 3 months. In total, this yielded individual-level estimates of Bolting under eight to nine distinct vernalization lengths for each population (Table S8A).

For each bolting estimate, we calculated the associated *Vernalization length* for early and late spring visits to each garden as the number of days with minimum air temperature < 4 °C (from dataloggers) from the day of transplant to the day of the visit. For the “no vernalization” cohort, vernalization length was set to zero. Following vernalization, it takes *C. americana* around a month to develop a characteristic bolting phenotype (Perrier *et al*., 2025). To account for the lag between receiving the required number of days of vernalization and the expression of cuing by bolting, we calculated the average number of days from the end of vernalization to bolting from previously published data for each population (*Bolting lag*, Table S8B, Perrier *et al*., 2025). This data was not available for four populations (KY1, KY5, MO2, OH1), so we imputed Bolting lag based on a linear model using the relationship between Bolting lag and latitude from the remaining populations (*R*^2^ = 0.36, *P* = 0.005, Fig. S3). We then subtracted the number of days with temperatures < 4 °C that fell within the lag period from the estimated Vernalization length of each population for each garden and visit. This yielded a conservative estimate of the Vernalization length that cued bolting in the gardens (Table S8B).

#### Geographic variation in vernalization requirement and sensitivity

We constructed reaction norms of Bolting against Vernalization length for each population. We modeled individual-level Bolting using a generalized linear mixed-effects model with a binomial error distribution and Vernalization length as a fixed effect using the R package *LmerTest* (Kuznetsova *et al*., 2017). Random effects included population, allowing both the intercept and the effect of Vernalization length to vary among populations (e.g. Fig. S4), as well as seed-family nested within population. We excluded the southern FL81 population from this and all subsequent analyzes as bolting was insensitive to Vernalization length due to consistently high bolting levels, including in plants grown without vernalization (Table S8A).

For each population, we used the reaction norms to estimate two metrics (Table S9A): V*ernalization requirement*, the number of days of vernalization at which half the individuals of a population are predicted to bolt (average bolting = 0.5; set to zero if a negative number of days), and *Vernalization sensitivity*, the slope of the Bolting–Vernalization length relationship on the logit scale. We then tested for geographic variation in these metrics in separate linear models assuming a gaussian distribution with latitude as a fixed effect. Preliminary analyses tested whether latitude was best described by a linear or quadratic relationship (Table S9B); results are reported for the best model.

### Q3: Does adaptation to past warming affect projected response to future climates?

Adaptations that likely allowed rear-edge populations to persist under past warming may improve their resilience to future climate change. Bolting was identified as the key to adaptation to *C. americana*’s warmest habitats (Q1, see Results), so we used this life stage to predict performance under future climates. We forecast bolting under future climates combining the bolting - vernalization reaction norms (Q2) with projections from multiple climate-change scenarios for each population in the common garden experiment. We then compare bolting forecasts among populations to test if rear-edge populations are more resilient or vulnerable than the rest of the range under expected future climates. Variables estimated for these analyses are summarized in Table S7B.

#### Estimates of future winter conditions

Climate datasets that allow the comparison of contemporary and future conditions do not provide the daily temperature information needed to estimate vernalization length. A suitable proxy may be the Minimum Temperature of the Coldest Month (BIO6), for which data under contemporary and predicted future climates is available from WorldClim 2.1 (Fick & Hijmans, 2017). To determine this, we estimated *Contemporary vernalization length* experienced by each population at their location of origin (days with minimum air temperature < 4 °C, PRISM temperature data, averaged 2018-2022, Table S10A). For each population’s location, we also obtained Minimum Temperature of the Coldest Month from WorldClim based on contemporary climates (1970-2020, 30-arc seconds resolution, Table S10A). These two estimates of winter cold are tightly associated (R^2^ = 0.96, BIO6 = 15.17 – (0.18 * vernalization length), Fig. S5A), indicating that Minimum Temperature of the Coldest Month is a suitable proxy for vernalization length.

We used this strong relationship to estimate *Future vernalization length* in the climate change scenarios estimated from the four Shared Socio-economic Pathways (SSP). We obtained projected Minimum Temperature of the Coldest Month values under SSP126, SSP245, SSP370 and SSP585 for 2081–2100 from the BCC-CSM2-MR climate model from WorldClim (30-arc seconds resolution, Table S10A). These scenarios represent a gradient of projected warming from mild to severe (estimated warming 1.8 °C, 2.7 °C, 3.6 °C and 4.4°C respectively, Intergovernmental Panel On Climate Change, 2023). We converted these estimates of future Minimum Temperature of the Coldest Month for the four scenarios into estimates of Future vernalization length for each population using the above equation (Table S10A, Fig. S5B).

#### Range-wide estimates of future bolting loss

We forecasted bolting loss, the reduction in bolting probability expected under projected future warmer winters relative to contemporary conditions, using the population-specific reaction norms developed above. For each population, we first predicted *Contemporary bolting probability* from the reaction norm (Table S10B) based on contemporary vernalization length estimated above (Table S10A). We then predicted *Future bolting probability*, again from the reaction norm (Table S10B), but using Future vernalization length at that location estimated for each climate change scenario (Table S10A). For each population and scenario, we then calculated *Bolting loss* as (bolting_future_ – bolting_contemporary_) / bolting_contemporary_ (Table S10B). We then tested for geographic variation in bolting loss using linear models with latitude as a fixed effect for each climate change scenario. These models included a quadratic latitude term to allow for potential non-linear relationships. Bolting loss was analyzed assuming a normal distribution.

#### Contribution of geographic variation in current and future winter climates

In addition to differences in reaction-norm shape, variation in bolting loss could, in part, be due to geographic variation in the magnitude of climate change. In particular, *C. americana* populations at higher latitudes are projected to experience a stronger reduction in future vernalization length than those at lower latitudes (Fig. S5B). To control for this variation, we created an estimate of bolting loss in which the magnitude of climate change was uniform across populations. To do this, we computed the *Expected change in vernalization length* as the difference between Contemporary and Future vernalization length for each population and climate change scenario. We then identified the maximum absolute difference across populations (Table S11A), and subtracted this value from the contemporary vernalization length of each population. This yielded an estimate of *Future vernalization length with fixed change* in vernalization length, where all populations are expected to experience the greatest warming predicted for any individual population (Table S11A). We then recalculated bolting loss under each climate scenario using Future vernalization length with fixed change to derive *Bolting loss with fixed change* (Table S11B) and tested for geographic variation as described above.

The difference between winter length and vernalization requirement is another potential source of variation in bolting loss among populations. Current winter length is longer than required to cue bolting in most populations (Table S11A), and likely provides them with a buffer in vernalization length before bolting is reduced by warming winters. However, populations from climates near their vernalization threshold are likely less buffered and may be susceptible to bolting loss even under a modest reduction in winter length. We created an estimate of bolting loss where the magnitude of the buffer is held fixed across populations to evaluate how variation in buffer length affects patterns of estimated bolting loss. To do this, we first estimated the *Vernalization buffer length* from the difference between the Contemporary vernalization length (Table S10A) and the Vernalization requirement (Table S9A). We then identified the shortest Vernalization buffer length across populations (AL21, Table S11A), and added this value to the Vernalization requirement of each population, yielding an estimate of *Contemporary vernalization length with fixed buffer,* where all populations experience the smallest buffer (Table S11A). We recalculated projected future vernalization length of each population by subtracting the Expected change in vernalization length under each scenario (estimated above) from the estimated Contemporary vernalization length with fixed buffer (Table S11A). We then used this *Future vernalization length with fixed buffer* to calculate *Contemporary bolting probability with fixed buffer*, *Future bolting probability with fixed* buffer, and *Bolting loss with fixed buffer* (Table S7B, S11C), and tested for geographic variation in Bolting loss with fixed buffer as described above.

## Results

### Q1: Which life stages are key for climate adaptation at the southern rear edge?

#### Which life stage shows the strongest local adaptation in southern rear-edge populations?

Out of the five life stages tested, we detected a significant relationship between life stage performance and ΔPC1, and significant differences in that relationship between gardens, only for survival over winter and bolting (Table 1, Fig. 1). Survival over winter had significant linear and quadratic relationships with ΔPC1 at the southern rear-edge (Table S12A), and the linear relationship was significantly stronger there than in the other two gardens (Table S12B, Fig.1B). However, this strong pattern was consistent with maladaptation in the southern rear-edge garden, with populations moved to the warmer southern rear-edge having greater survival over the winter than those native to that climate. Albeit not significant, a similar pattern of populations from colder climates having greater survival over winter and those from warmer climates smaller survival was found in the center garden (Fig.1B). Similarity between gardens suggests that variation in survival over winter may be an attribute of populations, independent of climate, and therefore does not reflect the genotype-environment interactions expected under local adaptation.

The only life stage with a strong signal of local adaptation was bolting, which was substantially greater for populations from similar climates in the southern rear-edge garden (significant ΔPC1, Table S12B). Probability of bolting for more northern populations was much smaller, including at or near zero (Fig.1C), consistent with local adaptation. In the other two gardens, bolting was uniformly high (Fig. 1C). In line with this result, local adaption was stronger at the southern rear-edge than in the other gardens (garden-ΔPC1 interaction, Table S12A, B, Fig. 1C). Finally, across all life stages in all gardens, bolting in the southern rear-edge garden had the greatest variation in performance among populations, with absolute differences in bolting proportion close to 1.

#### Which life stage contributes the most to local adaptation in the southern rear-edge?

Consistent with strong local adaptation for bolting, this was the only life stage that showed a strong and consistent relationship with lifetime fitness in the southern rear edge (Table 2, Fig. S6). In contrast, survival until reproduction and to a lesser extent reproductive output contributed most to lifetime fitness in the center garden, while survival over winter and reproductive output contributed the most to lifetime fitness in the northern rear-edge garden (Table 2, Fig. S6). This suggests that across the life stages measured in this experiment, bolting is the most important for local adaptation to the southern rear edge climates (Fig. 1).

**Table 2:** Contribution of life stages to lifetime fitness in the common gardens (CG)

| Life stage | CG - Center |  | CG - RearN |  | CG - RearS |  |
| --- | --- | --- | --- | --- | --- | --- |
| | Partial $R^2$ | $\Delta AICc$ | Partial $R^2$ | $\Delta AICc$ | Partial $R^2$ | $\Delta AICc$ |
| Survival over winter | 0.21 | 1.65 | **0.80** | **32.70** | 0.28 | 3.38 |
| Bolting | NA | NA | 0.15 | -0.61 | **0.88** | **44.25** |
| Survival until reproduction | **0.76** | **28.92** | 0.01 | -3.88 | 0.00 | -4.17 |
| Flowering | 0.30 | 4.49 | 0.05 | -3.06 | 0.02 | -3.69 |
| Reproductive output | **0.63** | **19.40** | **0.77** | **29.13** | 0.15 | -0.47 |

Lifetime fitness was tested in separate linear models for each garden with population means for all life stages included as continuous predictors. The contribution of each trait was assessed by computing partial $R^2$ , and the difference in AICc ( $\Delta AICc$ ) between the model without the trait and the full model. Traits with a partial $R^2 > 0.5$ and that lead to an increase in AICc when removed were considered the highest contributors (bold).

### Q2: What adaptations allow high performance in southern rear-edge conditions?

Southern rear-edge populations have lower vernalization requirements and are less sensitive to vernalization length than populations in the rest of the range. Bolting significantly decreased under shorter vernalization lengths (positive β, Table S13). This relationship depended on population, with southern populations less affected by shorter vernalization lengths than more northern populations (Fig. 2A). The vernalization length required to reach an average bolting of 0.5 had a strong positive relationship with population latitude, except in the most northern populations where vernalization requirements changed little with increased latitude, yielding an overall quadratic relationship (Table S9B, S14, Fig. 2B). Vernalization sensitivity also increased linearly with latitude, with bolting probability being relatively insensitive to changes in vernalization length in the most southern populations and very sensitive in the most northern populations (Table S9B, S14, Fig. 2A, C). Overall, these results indicate that southern rear-edge populations are able to bolt following shorter winters, and their bolting is less dependent on winter length than populations in the rest of the range.

**Figure 2:**
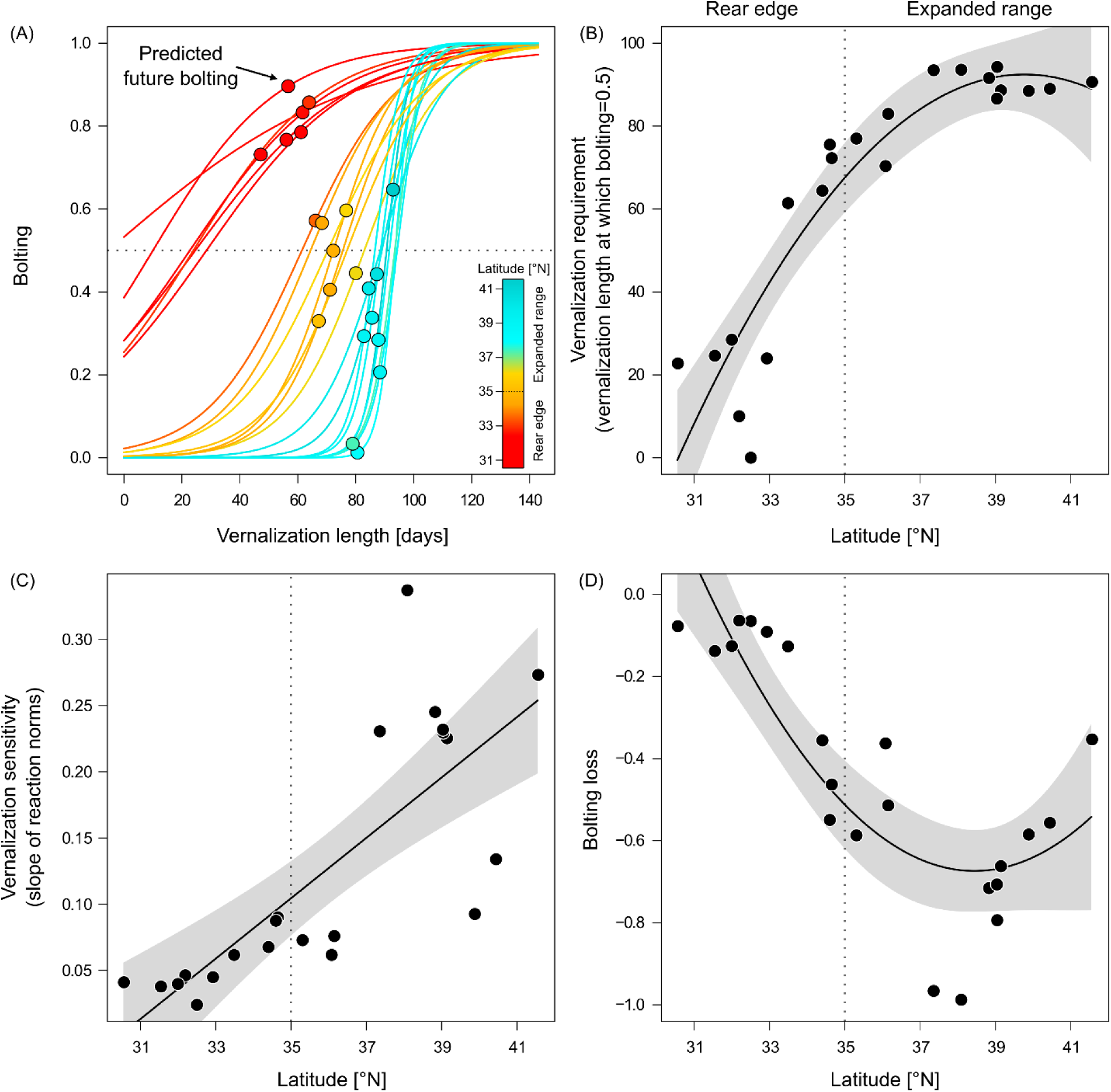
Reduced vernalization requirement and sensitivity make rear-edge populations resilient to climate change. **(A)** Model-predicted population-specific reaction norms (lines) of bolting across vernalization length (also Fig. S4). Colors distinguish populations by latitude. The horizontal dotted line indicates where predicted bolting crosses 0.5. Dots represent the estimation of Bolting probability under future climates (SSP 585, +4.4 °C) for each population. **(B)** Vernalization requirement, **(C)** Vernalization sensitivity and **(D)** Bolting loss under future climates (2081–2100, SSP 585) for each population (dots), estimated from the reaction norms in (A). The vertical dotted line indicates the separation between the rear edge (<35°N) and the expanded range. The solid line represents the significant relationship between each variable and latitude, with the 95% CI indicated in gray shading. Test statistics are reported in Table 3, S13 and S14.

**Table 3:** Variation in Bolting loss across the range for each climate change scenario.

| Scenario | $\beta$ latitude | $\beta$ latitude<sup>2</sup> | $F$ | | $R^2$ |
| --- | --- | --- | --- | --- | --- |
| SSP126 | -0.01 | 0.17 | 3.08 | (*) | 0.17 |
| SSP245 | -0.22 | 0.56 | **7.78** | ** | 0.39 |
| SSP370 | -0.53 | 0.68 | **11.37** | *** | 0.50 |
| SSP585 | -1.00 | 0.56 | **24.44** | *** | 0.69 |

Bolting loss under future climate was estimated for scenarios representing a range of future warming (SSP126 +1.8 °C, SSP245 +2.7 °C, SSP370 +3.6 °C, SSP585 +4.4 °C). Bolting loss was tested in separate linear models for each climate change scenario with latitude as a continuous quadratic predictor. Test statistics include the slope of the effect of latitude ( $\beta$ ) estimated for the linear and the squared term, the $F$ value (estimated jointly for both terms, bold if P-values < 0.05) and the $R^2$ of each model. (\*) $P < 0.1$ ; \* $P < 0.05$ ; \*\*\* $P < 0.001$ .

### Q3: Does adaptation to past warming affect projected response to future climates?

Southern rear-edge populations are expected to experience modest bolting loss (near zero) under projected future warming, even under the most severe warming scenario (Fig. 2D, S7). There was a positive quadratic relationship between bolting loss and latitude for each warming scenario (Table 3), with mid-latitude populations predicted to have larger reductions in bolting than low- and high-latitude populations (Fig. 2D, Fig. S7). Bolting loss was modest with little variation across populations under the mildest warming scenario (SSP126, Table 3, Fig. S7A), but was significant for the remaining scenarios (SSP245, SSP370, SSP585), increasing with the severity of predicted warming (Table 3, Fig. S7 B,C, Fig. 2D). Low- and high-latitude populations were predicted to experience similar modest bolting loss under the two intermediate warming scenarios (SSP245 Fig. S7B, SSP370 Fig. S7C). Under the strongest warming (SSP585), mid-latitude populations were predicted to experience up to 99% less bolting than in current climates (Fig. 2D). Under this scenario, high-latitude populations are also projected to experience strong reductions in bolting, with low-latitude populations experiencing little bolting loss (Fig. 2D).

Geographic patterns in projected bolting loss are not driven by variation in the magnitude of climate change among populations. When the change in vernalization length expected under future climates was held constant across latitudes, we observed a similar pattern of stronger bolting loss at mid-latitudes as described above (Table S15, Fig. 3A, Fig. S8). The main difference was that southern populations showed stronger reduction in bolting probability across climate change scenarios.

**Figure 3:**
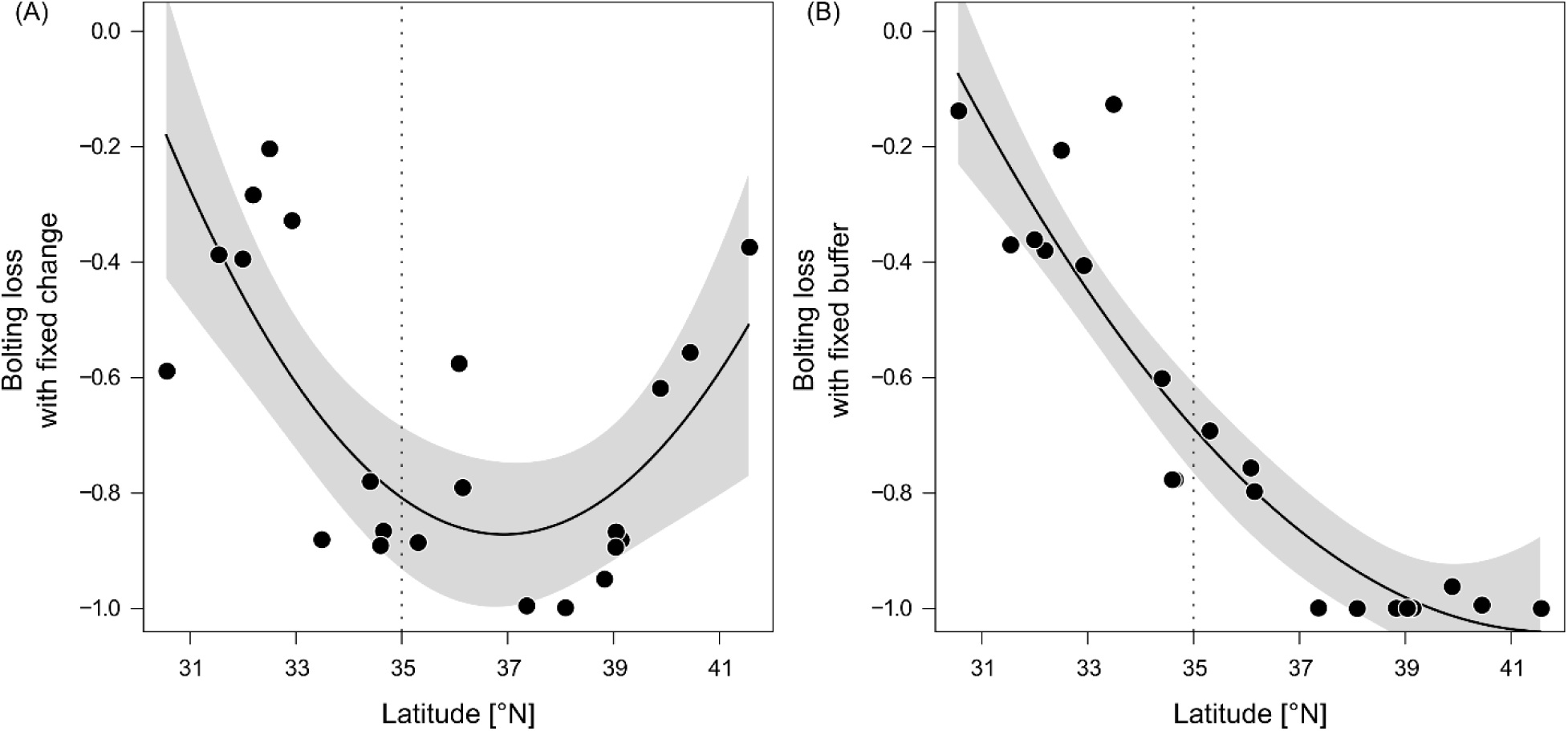
Rear-edge populations are more resilient even after accounting for the magnitude of climate change (A) and climate buffering (B). Bolting loss estimated under future climates (2081–2100, SSP 585, +4.4 °C) for each population (dots), with the expected change in vernalization length (A) and the length of the vernalization buffer (B) held constant across the range. The vertical dotted line indicates the separation between the rear edge (<35°N) and the expanded range. The solid line represents the significant relationship between each variable and latitude, with the 95% CI indicated in gray shading. Test statistics are reported in Table S15 and S16.

In contrast, climate buffering strongly affects geographic patterns in bolting loss. When holding the buffer between contemporary vernalization conditions and a population’s vernalization requirement constant, bolting loss showed an almost linear relationship with latitude, with high latitude populations showing greater loss in bolting while southern populations retained high bolting, rather than mid-latitude populations showing the most bolting loss (Table S16, Fig. 3B, Fig. S9).

## Discussion

### Shifts in developmental cuing linked to milder winters drive local adaptation at the southern rear edge

Bolting, the transition from somatic growth to reproduction, is the key life stage requiring adaptation for persistence at the southern rear edge in *C. americana*. In the southern garden, only bolting showed patterns consistent with local adaptation (Kawecki & Ebert, 2004; Leimu & Fischer, 2008). Bolting was high only for the most local populations, and declined to zero as climate mismatch increased. Bolting was also the most important determinant of lifetime fitness at the southern rear-edge garden. In contrast, bolting was uniformly high regardless of population origin in the center and northern rear-edge gardens, indicating that it doesn’t contribute to local adaptation outside the southern rear edge. Together, our results indicate that local adaptation to the subtropical southern rear-edge climates is driven by adaptation at a key life stage transition. Whether developmental transitions are common drivers of rear-edge adaptation, or specific to this species, is unclear. Studies of local adaptation at rear edges remain rare (Perrier et al., 2026b) and generally focus on a few targeted traits (Mathiasen & Premoli, 2016), especially those linked to heat stress response (Saada *et al*., 2016; King *et al*., 2019), or on contemporary evolution as demonstrated in resurrection studies (Anstett *et al*., 2026; Sheth *et al*., 2026). Our study highlights the importance of using explicit fitness-based tests across the life cycle to decipher rear-edge adaptation, and joins growing calls for more empirical work on adaptation at range limits (Angert *et al*., 2020; Bontrager *et al*., 2021).

Climates in rear-edge habitats are often unsuitable relative to those in the rest of the range (Vilà-Cabrera *et al*., 2019; Bontrager *et al.,* 2021), including in *C. americana* (Perrier *et al*., 2026a; c). Lack of suitability in rear-edge climates is typically associated with summer heat and drought stress (Cahill *et al*., 2014). In contrast, our finding that bolting is the key adaptation to the southern rear-edge in *C. americana* indicates that adaptation to warmer conditions primarily involves the evolution of developmental cues linked to milder winter temperatures. In this species, bolting only occurs after vernalization, the prolonged exposure to winter cold, with inadequate vernalization generally resulting in a failure to bolt (Perrier *et al*., 2025). Winters in the subtropical climate at the southern rear edge provide few days sufficiently cold to induce vernalization. Consistent with this climate, we found that southern rear-edge populations have reduced vernalization requirements and sensitivity. This indicates that adaptation to warmer rear-edge climates involves the decoupling of reproductive cuing from prolonged cold, allowing high levels of bolting despite limited chilling. One of the southernmost populations, FL81, routinely had high levels of bolting, even in the absence of vernalization, indicating that this evolutionary trajectory can culminate in the complete loss of this cue. Temperate plants often exhibit reduced vernalization requirements toward lower latitudes (e.g. Wesselingh *et al*., 1994; Dijk *et al*., 1997; Boudry *et al*., 2002; Stinchcombe *et al*., 2005; Jokela *et al*., 2015). Inadequate cueing may therefore represent a widespread but underappreciated climatic constraint associated with rear-edges.

Why does local adaptation at the southern rear edge of *C. americana* primarily involve adaptation at a single life stage? Across plants and animals, the complexity of local adaptation varies tremendously and can differ with environmental context (Wadgymar *et al*., 2022). The number of traits involved in local adaptation may in part reflect environmental complexity, with environments imposing multiple, interacting stressors requiring coordinated multi-trait adaptation, whereas adaptation in environments limiting fitness along a single axis may be reached through changes to a single trait (White & Butlin, 2021). In *C. americana*, although rear-edge environments differ substantially from central parts of the range, this divergence largely occurs along a climatic axis of temperature (Perrier *et al*., 2026a, 2025). While we only investigated this axis, our results indicate fitness in warmer conditions was primarily limited by vernalization length, a component of the temperature axis. Populations experiencing substantial climate mismatch maintained high performance across other studied stages of the life cycle indicating limited effects of the temperature axis on these fitness components (though notably germination was not tested). Therefore, rear-edge climates in *C. americana* appear to impose a specific, simple constraint on fitness, favoring adaptation at a critical life-history transition, rather than coordinated evolution across the life cycle.

Alternative explanations for the predominant role of bolting in adaptation to southern rear-edge habitats in *C. americana*, while possible, seem unlikely. For example, the cuing of reproduction through vernalization may be the only life stage with adequate genetic variation to respond to such strong climate selection. Bolting, indeed, exhibited the greatest phenotypic variation across populations in the southern rear-edge garden (Fig. 1). However, survival over winter also varied with climate difference, indicating sufficient variation for selection to act on. Another possibility is that bolting represents a severe fitness bottleneck such that individuals failing to bolt experience complete reproductive failure, regardless of their performance in later life stages. Strong selection on bolting could thus mask selection later in the life cycle (Grafen, 1988; Mittell & Morrissey, 2024). However, individuals of the summer cohort (adequate vernalization) raised in the southern rear-edge garden showed generally high post-bolting performance, suggesting that selection at later-life stages was not masked by this bottleneck, but instead was weak.

Our study provides insight into the processes that likely shaped rear-edge evolution under past climate change. Across species, rear edges have differed in their response to warming during the postglacial period. Some have persisted in the dramatically warmer habitats, showing strong local adaptation (Bontrager *et al*., 2021). Others have declined and even experienced local extinction (Hampe & Petit, 2005), potentially reflecting lags in adaptation to sustained postglacial warming (Burger & Lynch, 1995; Davis & Shaw, 2001; Chevin *et al*., 2010; Perrier *et al*., 2026b). Many factors may limit adaptation at rear edges, including a lack of standing genetic variation, genetic drift, or trade-offs (Antonovics, 1976; Polechová & Barton, 2015; Polechová, 2018; Willi & Van Buskirk, 2019, 2022; Perrier *et al*., 2026b). Our findings that a single life-stage is the main driver of rear-edge local adaptation in *C. americana* suggests that the persistence of refugial populations under climate change may in part depend on the complexity of environmental stress, and therefore the complexity of required adaptation. Adaptation involving one or a few traits may be more readily achieved than coordinated multi-trait adaptation, as simultaneous evolutionary change across multiple traits can be constrained by genetic correlations and incur higher selective costs (Antonovics, 1976; Orr, 2000; Bishop *et al*., 2026). Identifying whether successful adaptation to marginal rear-edge conditions is contingent on single-trait or multi-trait niche evolution is a crucial step in improving our understanding of the constraints on persistence under future climates at warmer range limits.

### Rear-edge responses to future warming and the legacy of past adaptation

Across temperate species, rear edges are generally expected to face elevated risks of population decline and extinction under future warming, as they are likely to experience conditions beyond species’ thermal niche limits (Thomas *et al*., 2004; Hampe & Petit, 2005; Nadeau & Urban, 2019; Lenoir *et al*., 2024). This may be particularly true for temperate plants that rely on cold cues to regulate their life cycle, where inadequate cuing under future warmer winters may lead to reproductive failure and associated fitness loss. While our projections indicate that all populations across *C. americana*’s range are likely to experience some reduction in bolting under future warming due to shorter winters, southern rear-edge populations are predicted to experience the smallest reduction, even under extreme warming scenarios. With bolting being a key component of fitness under warm climates in the species, our results indicate that *C. americana’s* southern rear-edge populations are the least likely to experience fitness loss under future climates. This limited vulnerability reflects a reduced dependence on winter cues that, as discussed above, likely evolved in response to past warming. Together, these findings join growing evidence that rear edges may not be as vulnerable to climate change as commonly thought (Vilà-Cabrera & Jump, 2019) and highlight how likely evolutionary responses to past warming can mitigate fitness loss under future climate change.

Multiple factors interact to determine vulnerability to future warmer winters resulting in non-linear patterns of predicted bolting loss across *C. americana*’s range. The modest bolting loss expected in southern rear-edge populations reflects the evolution of shorter vernalization requirements and reduced sensitivity to changes in vernalization, as well as a smaller expected reduction in winter conditions relative to populations elsewhere in the range. With this explanation, bolting loss might be expected to scale monotonically with latitude, as the vernalization requirement, sensitivity and the magnitude of climate change generally increase northward. However, the strongest loss was predicted for mid-latitude populations, with northern populations showing intermediate loss. This pattern results from mid-latitude populations occurring in climates that closely match their current vernalization requirement such that even modest reductions in winter chilling push them beyond a reproductive threshold. In contrast, northern populations appear to be buffered by contemporary winters that exceed their vernalization requirement, such that only the most extreme warming scenarios are predicted to result in inadequate vernalization.

Vulnerability to future climates may also be effected by adaptation at life stages other than bolting. In the southern rear-edge garden, some populations originating from much cooler areas appear to experience somewhat reduced survival until reproduction, flowering and reproductive output relative to local populations, which suggests that these populations could incur fitness loss under future warming beyond that expected from inadequate bolting. However, the decline in performance at these life stages was much weaker than the decline due to bolting, and not statistically supported. Life stages other than bolting may also affect southern rear-edge populations, as they had lower survival over winter in the southern rear-edge garden than populations at mid-latitudes. However, southern rear-edge populations had lower survival over winter than mid-latitude population across all gardens, suggesting it is an intrinsic property of the populations and may not worsen under future warming. In total, bolting loss is expected to be the dominant cause of fitness loss under future warming.

Rear-edge populations vary in their predicted response to future climates, with southern populations maintaining high predicted bolting rates while northern rear-edge populations (∼35N) likely experiencing substantial bolting loss. This variation may be linked to differences in evolutionary legacy among the populations, despite having a shared origin as refugial populations (Perrier *et al.,* 2026c). The southernmost populations experienced substantially stronger habitat decline under postglacial climate change (Perrier *et al.,* 2026c), and express stronger local adaptation (Perrier *et al*., 2026a) than the northern rear-edge populations. This suggests prolonged exposure to past warming may have been key in evolving adaptive strategies predicted to be beneficial under future warming. Such adaptation appears restricted to the genetically distinct southern rear edge (Perrier *et al.,* 2026c). The specific adaptation observed in this study, decoupling of reproductive transitions from environmental cues, may be particularly beneficial under future climates as it buffers populations against the disruption of ancestral cue–environment relationships (Anderson, 2016). Understanding the conditions under which adaptation was required for populations to persist under past warming, and the conditions under which predicted past adaptation can promote future resilience, may provide insight into why warm-edge populations persist under climate change in some species while declining in others.

Our findings have important implications for forecasting range dynamics under future climates. Species are commonly predicted to shift their distributions poleward and upward in elevation in response to warming, with range contractions at warm range limits (Lenoir *et al*., 2024; Urban, 2024). However, as we show here, populations at the warm edge of *C. americana* are projected to experience the least fitness decline under future warming, suggesting that future warm-edge contraction is unlikely or will be modest in this species. Instead, the extremely low bolting rates predicted for populations at mid-latitudes may lead to their decline and even loss, resulting in a range split rather than a range shift. Observational studies of species’ range shifts under ongoing warming increasingly find warm edges lagging behind expected contraction, particularly in plant species (Alexander *et al*., 2018; Geppert *et al*., 2020; Duchenne *et al*., 2021; Rubenstein *et al*., 2023). These expectations are often derived from ecological niche and species distribution models (Pacifici *et al*., 2015), which typically assume uniform fitness-environment relationships across species’ ranges (Wiens *et al*., 2009). Our results demonstrate that this assumption can be violated when populations differ in local adaptation. Models that ignore local adaptation and population-specific performance–climate relationships may systematically overestimate vulnerability at warm range limits and miss possible population decline in other parts of the range (Valladares *et al*., 2014; Benito Garzón *et al*., 2019; Habibzadeh *et al*., 2021). Incorporating local adaptation into forecasts is critical to accurately predict whether, and under what conditions, warm-edge populations will persist or decline under future climates.

## Conclusion

Understanding evolution at species’ warm range limits provides critical insight into the ecological and evolutionary processes shaping responses to past and future climate change (Brodie *et al*., 2025). Our findings show that persistence of rear-edge populations under past warming did not require complex multi-trait shifts or tolerance to extreme abiotic stress but instead likely involved adaptation in one key fitness-limiting life-history transition that permits persistence in the face of weakened environmental cues. More broadly, our study demonstrates that such presumed evolutionary legacies of past climate change can strongly influence the response of warm-edge populations to future warming. In *C. americana*, adaptation that likely enabled persistence under historical warming reduces vulnerability to future change, highlighting how alignment between past and future selective pressures can promote resilience at range limits. Under such conditions, rear-edge populations may harbor adaptive variation that is beneficial under warming climates, and may serve as a source of adaptive potential for other parts of the species’ range (Prober *et al*., 2015; Meek *et al*., 2023). Overall, our results emphasize that accounting for the evolutionary context of range-edge populations is essential for improving predictions of species’ range shifts.

## Acknowledgements

This work was supported by the Swiss National Science Foundation (P2BSP3_195363), the National Science Foundation (NSF DEB-2140189) and the University of Virginia College of Arts and Sciences. We are grateful to David Westneat (University of Kentucky Ecological Research and Education Center Field Station, Lexington, KY), Gordon Burghardt (Blaine, TN), Trevor Stamey (Clemson University Experimental Forest, Clemson, SC), Theresa Jepsen (Florida State University Mission Road Research Facility, Tallahassee, FL) for logistical support in establishing common gardens. For their help in raising of plants, performing crosses, and setting up the common garden experiments, we thank A. Burricks, C. Claussen, S. Cox, K. Haines, O. Keenan, S. Kelly, K. Lamb, A. López, H. Makowski, M. Marcich, L. Pizarro, E. Scott, E. Savier, & M. Turner. We are also grateful to J. Collins, L. Elliott, E. Galloway, J. Hansen, H. Horne, J. Kees, M. Kohout, R. Laporte, D. Reed and B. Sutherland for seed collection in natural populations. Collection permits were provided by the Florida Department of Environmental Protection, the Tennessee Department of Environment and Conservation, and Missouri Department of Conservation.

## Author Contributions

All authors contributed to the study design. AP collected seeds in the field, performed crosses, raised plants in the common garden experiment, and analyzed the data. AP wrote the manuscript with input from LG.

## Data availability statement

Fitness and climate data necessary to recapitulate the analyses presented in this study are stored on Zenodo (https://doi.org/10.5281/zenodo.23247681).

## References

Aguirre-Liguori, J. A., Ramírez-Barahona, S., & Gaut, B. S. (2021). The evolutionary genomics of species’ responses to climate change. Nature Ecology & Evolution, 5, 1350–1360.

Alexander, J. M., Chalmandrier, L., Lenoir, J., Burgess, T. I., Essl, F., Haider, S., Kueffer, C., McDougall, K., Milbau, A., Nuñez, M. A., Pauchard, A., Rabitsch, W., Rew, L. J., Sanders, N. J., & Pellissier, L. (2018). Lags in the response of mountain plant communities to climate change. Global Change Biology, 24, 563–579.

Anderson, J. T. (2016). Plant fitness in a rapidly changing world. New Phytologist, 210, 81–87.

Anderson, J. T., DeMarche, M. L., Denney, D. A., Breckheimer, I., Santangelo, J., & Wadgymar, S. M. (2025). Adaptation and gene flow are insufficient to rescue a montane plant under climate change. Science, 388, 525–531.

Angert, A. L., Bontrager, M. G. & Ågren, J. (2020). What do we really know about adaptation at range edges? Annual Review of Ecology, Evolution, and Systematics, 51, 341–361.

Anstett, D. N., Anstett, J., Sheth, S. N., Moxley, D. R., Branch, H. A., Jahani, M., Huang., K, Todesco, M., Jordan, R., Lazaro-Guevara, J. M., Rieseberg, L. H., & Angert, A. L. (2026). Rapid evolution predicts demographic recovery after extreme drought. Science, 391, 1172–1176

Antonovics, J. (1976). The nature of limits to natural selection. Annals of the Missouri Botanical Garden, 63, 224.

Assis, J., Araújo, M. B., & Serrão, E. A. (2018). Projected climate changes threaten ancient refugia of kelp forests in the North Atlantic. Global Change Biology, 24, e55–e66.

Barnard-Kubow, K. B., Debban, C. L., & Galloway, L. F. (2015). Multiple glacial refugia lead to genetic structuring and the potential for reproductive isolation in a herbaceous plant. American Journal of Botany, 102, 1842–1853.

Baskin, J. M., & Baskin, C. C. (1984). The ecological life cycle of *Campanula americana* in Northcentral Kentucky. Bulletin of the Torrey Botanical Club, 111, 329–337.

Bates, D., M. Mächler, B. Bolker, & Walker, S. 2015. Fitting linear mixed-effects models using lme4. Journal of Statistical Software 67:1–48. 10.18637/jss.v067.i01

Benito Garzón, M., Robson, T. M., & Hampe, A. (2019). ΔTraitSDMs: Species distribution models that account for local adaptation and phenotypic plasticity. New Phytologist, 222, 1757–1765.

Bishop, S. G. D, Stinchcombe, J. R., & Baucom, R. S. (2026). A resurrection experiment reveals reduced adaptive potential in a common agricultural weed, Evolution Letters, 10, 451–462.

Bontrager, M., Usui, T., Lee-Yaw, J. A., Anstett, D. N., Branch, H. A., Hargreaves, A. L., Muir, C. D., & Angert, A. L. (2021). Adaptation across geographic ranges is consistent with strong selection in marginal climates and legacies of range expansion. Evolution, 75, 1316–1333.

Boudry, P., Mccombie, H., & Van Dijk, H. (2002). Vernalization requirement of wild beet *Beta vulgaris* ssp. *maritima*: Among population variation and its adaptive significance. Journal of Ecology, 90, 693–703.

Brodie, J. F., Freeman, B. G., Mannion, P. D., & Hargreaves, A. L. (2025). Shifting, expanding, or contracting? Range movement consequences for biodiversity. Trends in Ecology & Evolution, 40, 439–448.

Burger, R. & Lynch, M. 1995. Evolution and extinction in a changing environment: a quantitative-genetic analysis. Evolution, 49, 151–163.

Cahill, A. E., Aiello-Lammens, M. E., Caitlin Fisher-Reid, M., Hua, X., Karanewsky, C. J., Ryu, H. Y., Sbeglia, G. C., Spagnolo, F., Waldron, J. B., & Wiens, J. J. (2014). Causes of warm-edge range limits: Systematic review, proximate factors and implications for climate change. Journal of Biogeography, 41, 429–442.

Chevin, L.-M., Lande, R. & Mace, G.M. 2010. Adaptation, plasticity, and extinction in a changing environment: towards a predictive theory. PLOS Biology, 8, e1000357.

Cinelli, C., & Hazlett, C. (2020). Making sense of sensitivity: Extending omitted variable bias. Journal of the Royal Statistical Society Series B: Statistical Methodology, 82, 39–67.

Davis, M. B., & Shaw, R. G. (2001). Range shifts and adaptive responses to Quaternary climate change. Science, 292, 673–679.

de Lafontaine, G., Napier, J. D., Petit, R. J., & Hu, F. S. (2018). Invoking adaptation to decipher the genetic legacy of past climate change. Ecology, 99, 1530–1546.

DeMarche, M. L., Doak, D. F., & Morris, W. F. (2018). Both life-history plasticity and local adaptation will shape range-wide responses to climate warming in the tundra plant *Silene acaulis*. Global Change Biology, 24, 1614–1625.

Dijk, H. V., Boudry, P., McCombre, H., & Vernet, P. (1997). Flowering time in wild beet (*Beta vulgaris* ssp. Maritima) along a latitudinal cline. Acta Oecologica, 18, 47–60.

Dorie, V., Bates, D., M. Mächler, B. Bolker, & Walker, S. (2026), blme: Bayesian Linear Mixed-Effects Models. R package version 1.0-7. Available at: http://CRAN.R-project.org/package=blme (accessed 19 May 2026).

Duchenne, F., Martin, G., & Porcher, E. (2021). European plants lagging behind climate change pay a climatic debt in the North, but are favoured in the South. Ecology Letters, 24, 1178–1186.

Fick, S. E., & Hijmans, R. J. (2017). WorldClim 2: New 1-km spatial resolution climate surfaces for global land areas. International Journal of Climatology, 37, 4302–4315.

Franks, S.J., Weber, J.J. & Aitken, S.N. 2014. Evolutionary and plastic responses to climate change in terrestrial plant populations. Evolutionary Applications 7: 123–139.

Galloway, L. F. (2002). The effect of maternal phenology on offspring characters in the herbaceous plant Campanula americana. Journal of Ecology, 90, 851–858.

Garnier, E., Navas, M.-L., & Grigulis, K. (2015). Gradients, response traits, and ecological strategies. In E. Garnier, M.-L. Navas, & K. Grigulis (Eds.), Plant Functional Diversity: Organism traits, community structure, and ecosystem properties. Oxford University Press.

Geppert, C., Perazza, G., Wilson, R. J., Bertolli, A., Prosser, F., Melchiori, G., & Marini, L. (2020). Consistent population declines but idiosyncratic range shifts in Alpine orchids under global change. Nature Communications, 11, 5835.

Ghouil, H., Sancho-Knapik, D., Ben Mna, A., Amimi, N., Ammari, Y., Escribano, R., Alonso-Forn, D., Ferrio, J. P., Peguero-Pina, J. J., & Gil-Pelegrín, E. (2020). Southeastern rear edge populations of *Quercus suber* L. showed two alternative strategies to cope with water stress. Forests, 11, 1344

Grafen, A. 1988. On the Uses of Data on Lifetime Reproductive Success. Chicago, IL: University of Chicago Press.

Habibzadeh, N., Ghoddousi, A., Bleyhl, B., & Kuemmerle, T. (2021). Rear-edge populations are important for understanding climate change risk and adaptation potential of threatened species. Conservation Science and Practice, 3, e375.

Hamann, E., Kesselring, H. & Stöcklin, J. 2018. Plant responses to simulated warming and drought: a comparative study of functional plasticity between congeneric mid and high elevation species. Journal of Plant Ecology, 11, 364–374.

Hampe, A., & Petit, R. J. (2005). Conserving biodiversity under climate change: The rear edge matters. Ecology Letters, 8, 461–467.

Hargreaves, A. L., Germain, R. M., Bontrager, M., Persi, J. & Angert, A.L. 2020. Local adaptation to biotic interactions: A Meta-analysis across Latitudes. The American Naturalist, 195, 395– 411.

Hewitt, G. (2000). The genetic legacy of the Quaternary ice ages. Nature, 405, 907–913.

Hewitt, G. M. (2004). Genetic consequences of climatic oscillations in the Quaternary. Philosophical Transactions of the Royal Society of London. Series B: Biological Sciences, 359, 183–195.

Hughes, P. D., Gibbard, P. L., & Ehlers, J. (2013). Timing of glaciation during the last glacial cycle: Evaluating the concept of a global ‘Last Glacial Maximum’ (LGM). Earth-Science Reviews, 125, 171–198.

Intergovernmental Panel On Climate Change (2023). Climate Change 2021 – The Physical Science Basis: Working Group I Contribution to the Sixth Assessment Report of the Intergovernmental Panel on Climate Change (1st ed.). Cambridge University Press.

Jackson, J., Le Coeur, C., & Jones, O. (2022). Life history predicts global population responses to the weather in terrestrial mammals. eLife, 11, e74161.

Jokela, V., Trevaskis, B., & Seppänen, M. M. (2015). Genetic variation in the flowering and yield formation of timothy (*Phleum pratense* L.) accessions after different photoperiod and vernalization treatments. Frontiers in Plant Science, 6.

Kawecki, T. J., & Ebert, D. (2004). Conceptual issues in local adaptation. Ecology Letters, 7, 1225–1241.

King, N.G., McKeown, N.J., Smale, D.A., Wilcockson, D.C., Hoelters, L., Groves, E.A., et al. 2019. Evidence for different thermal ecotypes in range centre and trailing edge kelp populations. Journal of Experimental Marine Biology and Ecology, 514-515: 10–17.

Kooyers, N.J. 2015. The evolution of drought escape and avoidance in natural herbaceous populations. Plant Science, 234, 155–162.

Koski, M. H., Layman, N. C., Prior, C. J., Busch, J. W., & Galloway, L. F. (2019). Selfing ability and drift load evolve with range expansion. Evolution Letters, 3, 500–512.

Kristensen, T. N., Ketola, T., & Kronholm, I. (2020). Adaptation to environmental stress at different timescales. Annals of the New York Academy of Sciences, 1476, 5–22.

Kuznetsova, A., Brockhoff, P. B., & Christensen, R. H. B. (2017). lmerTest Package: Tests in Linear Mixed Effects Models. Journal of Statistical Software, 82, 1–26.

Landoni, B., Suárez-Montes, S., Habeahan, R. H. F., Brennan, A. C., & Pérez-Barrales, R. (2024). Local climate and vernalization sensitivity predict the latitudinal patterns of flowering onset in the crop wild relative *Linum bienne* Mill. Annals of Botany, 134, 117–130.

Lamb, K., Debban, C. L., & Galloway, L. F. (2024). Phylogeography and paleoclimatic range dynamics explain variable outcomes to contact across a species’ range. Molecular Ecology, 33, e17450.

Leimu, R., & Fischer, M. (2008). A meta-analysis of local adaptation in plants. PLOS ONE, 3, e4010.

Lenoir, J., & Svenning, J.-C. (2015). Climate-related range shifts – a global multidimensional synthesis and new research directions. Ecography, 38, 15–28.

Lenoir, J., Svenning, J.-C., & Sheffer, M. (2024). Latitudinal and Elevational Range Shifts Under Contemporary Climate Change. In: Encyclopedia of Biodiversity, 690–709

Lenth, R. V., Piaskowski, J., Banfai, B., Bolker, B., Buerkner, P., Giné-Vázquez, I., Hervé, M., Jung, M., Love, J., Miguez, F., Riebl, H., & Singmann, H. (2025). emmeans: Estimated Marginal Means, aka Least-Squares Means (Version 2.0.1). https://cran.r-project.org/web/packages/emmeans/index.html

Mathiasen, P., & Premoli, A. C. (2016). Living on the edge: Adaptive and plastic responses of the tree *Nothofagus pumilio* to a long-term transplant experiment predict rear-edge upward expansion. Oecologia, 181, 607–619.

Meek, M. H., Beever, E. A., Barbosa, S., Fitzpatrick, S. W., Fletcher, N. K., Mittan-Moreau, C. S., Reid, B. N., Campbell-Staton, S. C., Green, N. F., & Hellmann, J. J. (2023). Understanding local adaptation to prepare populations for climate change. BioScience, 73, 36–47.

Mittell, E. A., & Morrissey, M. B. (2024). The missing fraction problem as an episodes of selection problem. Evolution, 78, 601–611.

Nadeau, C. P., & Urban, M. C. (2019). Eco-evolution on the edge during climate change. Ecography, 42, 1280–1297.

Nicotra, A.B., Atkin, O.K., Bonser, S.P., Davidson, A.M., Finnegan, E.J., Mathesius, U., Poot, P., Purugganan, M. D., Richards, C. L., Valladeres, F., & van Kleunen, M. (2010). Plant phenotypic plasticity in a changing climate. Trends in Plant Science 15: 684–692.

Orr, H. A. (2000). Adaptation and the cost of complexity. Evolution, 54, 13–20.

Pacifici, M., Foden, W. B., Visconti, P., Watson, J. E. M., Butchart, S. H. M., Kovacs, K. M., Scheffers, B. R., Hole, D. G., Martin, T. G., Akçakaya, H. R., Corlett, R. T., Huntley, B., Bickford, D., Carr, J. A., Hoffmann, A. A., Midgley, G. F., Pearce-Kelly, P., Pearson, R. G., Williams, S. E., Willis, S. G., Young, B., & Rondinini, C. (2015). Assessing species vulnerability to climate change. Nature Climate Change, 5, 215–224.

Pacifici, M., Visconti, P., Butchart, S. H. M., Watson, J. E. M., Cassola, F. M., & Rondinini, C. (2017). Species’ traits influenced their response to recent climate change. Nature Climate Change, 7, 205–208.

Parmesan, C., & Yohe, G. (2003). A globally coherent fingerprint of climate change impacts across natural systems. Nature, 421, 37–42.

Pelletier, E., & de Lafontaine, G. (2023). Jack pine of all trades: Deciphering intraspecific variability of a key adaptive trait at the rear edge of a widespread fire-embracing North American conifer. American Journal of Botany, 110, e16111.

Perrier A., Keenan O. J., Busch J. W., & Galloway L. F. (2026a). The legacy of past climate warming: strong local adaptation in rear-edge populations. Evolution letters, 10, 343–354.

Perrier A., Keenan, O. J., & Galloway L. F. (2026b). Revisiting evolution at the rear edge. Trends in Ecology and Evolution, 41, 439–450.

Perrier A., Lamb KC. & Galloway, L.F. (2026c). Heterogeneous evolutionary history defines the rear edge of the North American herb *Campanula americana*. Journal of Biogeography, 53, e70226.

Perrier, A., Turner, M. C., & Galloway, L. F. (2025). Shifts in vernalization and phenology at the rear edge hold insight into the adaptation of temperate plants to future milder winters. New Phytologist, 246, 1377–1389.

Polechová, J. 2018. Is the sky the limit? On the expansion threshold of a species’ range. PLoS Biology, 16, e2005372.

Polechová, J. & Barton, N.H. 2015. Limits to adaptation along environmental gradients. Proceedings of the National Academy of Sciences, 112, 6401–6406.

Prior, C. J., Layman, N. C., Koski, M. H., Galloway, L. F., & Busch, J. W. (2020). Westward range expansion from middle latitudes explains the Mississippi River discontinuity in a forest herb of eastern North America. Molecular Ecology, 29, 4473–4486.

Prober, S., Byrne, M., McLean, E., Steane, D., Potts, B., Vaillancourt, R., & Stock, W. (2015). Climate-adjusted provenancing: A strategy for climate-resilient ecological restoration. Frontiers in Ecology and Evolution, 3, 65.

R Core Team. (2026). R: a language and environment for statistical computing. R Foundation for Statistical Computing.

Rubenstein, M. A., Weiskopf, S. R., Bertrand, R., Carter, S. L., Comte, L., Eaton, M. J., Johnson, C. G., Lenoir, J., Lynch, A. J., Miller, B. W., Morelli, T. L., Rodriguez, M. A., Terando, A., & Thompson, L. M. (2023). Climate change and the global redistribution of biodiversity: Substantial variation in empirical support for expected range shifts. Environmental Evidence, 12, 7.

Saada, G., Nicastro, K. R., Jacinto, R., McQuaid, C. D., Serrão, E. A., Pearson, G. A., & Zardi, G. I. (2016). Taking the heat: Distinct vulnerability to thermal stress of central and threatened peripheral lineages of a marine macroalga. Diversity and Distributions, 22, 1060–1068.

Sánchez-Salguero, R., Camarero, J. J., Gutiérrez, E., González Rouco, F., Gazol, A., Sangüesa-Barreda, G., Andreu-Hayles, L., Linares, J. C., & Seftigen, K. (2017). Assessing forest vulnerability to climate warming using a process-based model of tree growth: Bad prospects for rear-edges. Global Change Biology, 23, 2705–2719.

Sheth, S.N., Albano, L.J., Blanchard, C., Cook, E., Diaz, R., Gomez-Vega, X., Kutella, K., Moazed, M., Patel, M., Prange, J., Ramadoss, N., Regan, A., Riley, A., Hernandez, M. R., Rojas, J., Strebler, M., Verma, A., Villano, L., Waits, J., Wang, D., Wilborn-Pilotte, O., Diez, J., Flores-Rentería, L., Sexton, J. P., & Muir, C. D. (2026). Evolutionary responses to historic drought across the range of scarlet monkeyflower. The American Naturalist, 207, 135–155.

Stinchcombe, J. R., Caicedo, A. L., Hopkins, R., Mays, C., Boyd, E. W., Purugganan, M. D., & Schmitt, J. (2005). Vernalization sensitivity in *Arabidopsis thaliana* (Brassicaceae): The effects of latitude and FLC variation. American Journal of Botany, 92, 1701–1707.

Thomas, C. D., Cameron, A., Green, R. E., Bakkenes, M., Beaumont, L. J., Collingham, Y. C., Erasmus, B. F. N., de Siqueira, M. F., Grainger, A., Hannah, L., Hughes, L., Huntley, B., van Jaarsveld, A. S., Midgley, G. F., Miles, L., Ortega-Huerta, M. A., Townsend Peterson, A., Phillips, O. L., & Williams, S. E. (2004). Extinction risk from climate change. Nature, 427, 145–148.

Urban, M. C. (2024). Climate change extinctions. Science, 386, 1123–1128.

Valladares, F., Matesanz, S., Guilhaumon, F., Araújo, M. B., Balaguer, L., Benito-Garzón, M., Cornwell, W., Gianoli, E., van Kleunen, M., Naya, D. E., Nicotra, A. B., Poorter, H., & Zavala, M. A. (2014). The effects of phenotypic plasticity and local adaptation on forecasts of species range shifts under climate change. Ecology Letters, 17, 1351–1364.

Vilà-Cabrera, A., & Jump, A. S. (2019). Greater growth stability of trees in marginal habitats suggests a patchy pattern of population loss and retention in response to increased drought at the rear edge. Ecology Letters, 22, 1439–1448.

Vilà-Cabrera, A., Premoli, A. C., & Jump, A. S. (2019). Refining predictions of population decline at species’ rear edges. Global Change Biology, 25 1549–1560.

Wadgymar, S. M., DeMarche, M. L., Josephs, E. B., Sheth, S. N., & Anderson, J. T. (2022). Local adaptation: Causal agents of selection and adaptive trait divergence. Annual Review of Ecology, Evolution, and Systematics, 53, 87–111.

Wesselingh, R. A., Klinkhamer, P. G. L., Jong, T. J., & Schlatmann, E. G. M. (1994). A latitudinal cline in vernalization requirement in *Cirsium vulgare*. Ecography, 17, 272–277.

White, N. J., & Butlin, R. K. (2021). Multidimensional divergent selection, local adaptation, and speciation. Evolution, 75, 2167–2178.

Wiens, J. A., Stralberg, D., Jongsomjit, D., Howell, C. A., & Snyder, M. A. (2009). Niches, models, and climate change: Assessing the assumptions and uncertainties. Proceedings of the National Academy of Sciences, 106, 19729–19736.

Willi, Y., & Van Buskirk, J. (2019). A practical guide to the study of distribution limits. The American Naturalist, 193, 773–785.

Willi, Y., & Van Buskirk, J. (2022). A review on trade-offs at the warm and cold ends of geographical distributions. Philosophical Transactions of the Royal Society B: Biological Sciences, 377, 20210022.

