## Supplementary information for "Adaptation to marginal habitats at the rear edge provides resilience to future warming"

**Supplementary tables**

**Table S1: Populations used and replication in each cohort and common garden (CG) – Q1**

| | | | | Summer cohort (2023) | | | | | | Winter cohort (2023 – 2024) | | | | | | | |
|---|---|---|---|---|---|---|---|---|---|---|---|---|---|---|---|---|---|
| | | | | Families | | | Individuals | | | Families | | | | Individuals | | | |
| ID | Latitude [° N] | Longitude [° E] | Seed source | CG-Center | CG-RearN | CG-RearS | CG-Center | CG-RearN | CG-RearS | CG-Center | CG-Int | CG-RearN | CG-RearS | CG-Center | CG-Int | CG-RearN | CG-RearS |
| FL83 | 30.56468 | -84.95982 | Crosses | 13 | 15 | 15 | 20 | 29 | 30 | 15 | 15 | 15 | 15 | 29 | 29 | 28 | 26 |
| FL81 | 30.81165 | -85.22535 | Crosses | 14 | 15 | 15 | 20 | 30 | 29 | 15 | 15 | 15 | 15 | 29 | 29 | 30 | 30 |
| AL2 | 31.54793 | -87.51453 | Crosses | 15 | 15 | 15 | 20 | 30 | 30 | 14 | 15 | 15 | 15 | 27 | 28 | 28 | 28 |
| MS6 | 31.99803 | -89.35603 | Crosses | 13 | 15 | 15 | 20 | 30 | 30 | 15 | 15 | 15 | 15 | 28 | 28 | 30 | 29 |
| AL23 | 32.19395 | -86.78484 | Crosses | 15 | 15 | 15 | 20 | 30 | 29 | 15 | 15 | 15 | 15 | 29 | 29 | 29 | 29 |
| AL22 | 32.50376 | -87.50489 | Crosses | 13 | 15 | 15 | 20 | 30 | 29 | 15 | 15 | 15 | 15 | 29 | 30 | 29 | 29 |
| AL79 | 32.92932 | -88.20820 | Crosses | 13 | 15 | 15 | 19 | 30 | 30 | 15 | 14 | 15 | 13 | 29 | 28 | 28 | 25 |
| AL21 | 33.49109 | -86.79829 | Crosses | 13 | 15 | 15 | 20 | 30 | 29 | 15 | 15 | 15 | 15 | 29 | 29 | 28 | 28 |
| MS8 | 34.40427 | -88.83093 | Crosses | 13 | 15 | 14 | 20 | 29 | 29 | 15 | 15 | 15 | 15 | 30 | 30 | 30 | 30 |
| GA1 | 34.60074 | -84.69664 | Crosses | 14 | 15 | 15 | 19 | 30 | 30 | 15 | 15 | 14 | 14 | 28 | 29 | 26 | 25 |
| ALBG | 34.65346 | -86.51638 | Crosses | 13 | 15 | 15 | 19 | 30 | 29 | 13 | 15 | 15 | 15 | 25 | 29 | 29 | 30 |
| TN3 | 35.30932 | -90.06770 | Crosses | 13 | 15 | 15 | 20 | 30 | 29 | 15 | 15 | 15 | 15 | 29 | 29 | 29 | 30 |
| TN34 | 36.08222 | -86.29611 | Crosses | 15 | 15 | 15 | 20 | 30 | 30 | 15 | 15 | 15 | 15 | 30 | 30 | 30 | 30 |
| AR2 | 36.15235 | -94.30465 | Crosses | 13 | 15 | 15 | 20 | 29 | 30 | 15 | 15 | 15 | 15 | 29 | 30 | 30 | 30 |
| KY5 | 37.36005 | -84.77186 | Field | 14 | 15 | 15 | 20 | 29 | 30 | 15 | 15 | 15 | 15 | 29 | 30 | 28 | 30 |
| KY1 | 38.09215 | -84.98688 | Field | 11 | 15 | 15 | 19 | 29 | 30 | 13 | 13 | 14 | 12 | 26 | 25 | 26 | 22 |
| MO2 | 38.83025 | -92.28509 | Field | 12 | 14 | 14 | 18 | 27 | 29 | 14 | 14 | 14 | 13 | 27 | 28 | 26 | 26 |
| OH1 | 39.03935 | -84.32774 | Field | 12 | 15 | 15 | 20 | 29 | 27 | 15 | 15 | 15 | 15 | 30 | 29 | 30 | 30 |
| KS60 | 39.04742 | -95.68152 | Crosses | 14 | 15 | 15 | 20 | 30 | 29 | 15 | 15 | 15 | 15 | 30 | 30 | 30 | 28 |
| IN5 | 39.14583 | -86.54833 | Crosses | 13 | 15 | 15 | 19 | 30 | 29 | 15 | 15 | 15 | 14 | 29 | 28 | 30 | 26 |
| OH119 | 39.88500 | -83.99700 | Crosses | 13 | 15 | 15 | 20 | 30 | 29 | 15 | 15 | 15 | 15 | 28 | 28 | 29 | 27 |
| IN4 | 40.44806 | -86.93444 | Crosses | 13 | 15 | 15 | 19 | 30 | 29 | 15 | 15 | 15 | 15 | 29 | 30 | 29 | 29 |
| IA19 | 41.56611 | -90.47500 | Crosses | 13 | 15 | 15 | 19 | 29 | 30 | 14 | 14 | 15 | 14 | 27 | 28 | 30 | 28 |

Families and individuals depict the number of unique seed families and individuals per population that were transplanted in each garden and tracked for performance. Only pots with successful and timely germination were transplanted and tracked (96% of pots). In CG-Center in the summer cohort, the first 10 blocks had to be excluded due to a mudslide. Individuals raised in GC-Int were entirely excluded from the local adaptation study (Q1) due to premature mortality in the summer cohort, but individuals of the winter cohort were included in the bolting studies (Q2 and Q3). In total, 3768 seedlings were tracked over the two cohorts in the local adaptation study (Q1), and 2625 individuals were tracked in the winter cohort for the bolting studies (Q2 and 3).

**Table S2A: Location of common gardens (CG) and schedule of the summer cohort – Q1**

| | Latitude | Longitude | Transplant | Visit 1 | Visit 2 | Visit 3 |
|---|---|---|---|---|---|---|
| CG-Center | 38.08079 | -84.47134 | 08/04/2023 | 16/07/2023 | 12/08/2023 | 09/09/2023 |
| CG-Int | 36.18525 | -83.69820 | - | - | - | - |
| CG-RearN | 34.68746 | -82.87249 | 22/03/2023 | 14/07/2023 | 13/08/2023 | 06/09/2023 |
| CG-RearS | 30.45723 | -84.33319 | 02/03/2023 | 12/07/2023 | 17/08/2023 | - |

Individuals raised in GC-Int were excluded from the local adaptation study (Q1) due to premature mortality. For CG-Rear, 98% of plants were considered mature at the second visit, thus all plants were collected, and the garden was not visited a third time.

**TableS2B: Schedule of the winter cohort – Q1**

| | Transplant | Visit 1 early spring | Visit 2 late spring |
|---|---|---|---|
| CG-Center | 29/10/2023 | 28/03/2024 | 05/06/2024 |
| CG-Int | 04/11/2023 | 28/03/2024 | 04/06/2024 |
| CG-RearN | 12/11/2023 | 25/03/2024 | 31/05/2024 |
| CG-RearS | 14/11/2023 | 26/03/2024 | 02/06/2024 |

**Table S3A: Description of climatic variables – Q1**

| Variables | Description |
|---|---|
| *Seasonal climatic variables calculated for winter, spring and summer* | |
| Tmean | Seasonal average of daily mean temperatures |
| Tmin | Seasonal average of daily minimum temperatures |
| Tmax | Seasonal average of daily maximum temperatures |
| PPTsum | Seasonal sum of daily precipitation |
| *Seasonal climatic variables calculated for winter only* | |
| DayFrost | Number of days during winter where Tmin < 0°C |
| DayVerna | Number of days during winter where Tmean < 4°C |
| *Bioclimatic variables* | |
| Bio 2 | Mean diurnal range (mean of monthly (max. temp. – min. temp.)) |
| Bio 3 | Isothermality ((Bio 2 / Bio 7) ×100) |
| Bio 4 | Temp. seasonality (standard deviation of monthly mean temp. ×100) |
| Bio 7 | Temp. annual range (max. temp. of warmest month – min. temp. of coldest month) |
| Bio 15 | Precipitation seasonality (coefficient of variation of monthly total precipitation) |

**Table S3B: Climate variables and the first PC axis for populations and common gardens – Q1**

| | Winter | | | | | | Spring | | | | Summer | | | | | | | | | |
|---|---|---|---|---|---|---|---|---|---|---|---|---|---|---|---|---|---|---|---|---|
| | Tmean [° C] | Tmin [° C] | Tmax [° C] | PPTsum [mm] | DayFrost [days] | DayVerna [days] | Tmean [° C] | Tmin [° C] | Tmax [° C] | PPTsum [mm] | Tmean [° C] | Tmin [° C] | Tmax [° C] | PPTsum [mm] | bio2 [° C] | bio3 | bio4 [° C] | bio7 [° C] | bio15 [mm] | PC1 |
| *Populations* | | | | | | | | | | | | | | | | | | | | |
| FL83 | 13.93 | 7.63 | 20.22 | 524.69 | 15.00 | 3.00 | 21.69 | 15.06 | 28.31 | 154.39 | 27.47 | 22.04 | 32.90 | 542.59 | 12.03 | 41.56 | 628.11 | 28.94 | 28.03 | 5.83 |
| FL81 | 13.83 | 7.69 | 19.97 | 471.12 | 12.80 | 3.00 | 21.70 | 15.42 | 27.98 | 180.03 | 27.59 | 22.40 | 32.77 | 521.06 | 11.68 | 40.79 | 636.11 | 28.62 | 26.86 | 5.76 |
| AL2 | 12.05 | 5.26 | 18.84 | 484.74 | 36.40 | 9.00 | 20.01 | 12.70 | 27.33 | 177.67 | 26.63 | 20.99 | 32.28 | 458.20 | 12.99 | 42.40 | 675.34 | 30.64 | 26.79 | 4.22 |
| MS6 | 11.22 | 4.80 | 17.63 | 670.41 | 36.40 | 12.20 | 19.57 | 12.85 | 26.29 | 263.80 | 26.63 | 21.05 | 32.21 | 415.68 | 12.41 | 39.60 | 712.46 | 31.35 | 30.49 | 3.78 |
| AL23 | 11.30 | 4.92 | 17.68 | 577.98 | 36.20 | 11.60 | 19.54 | 12.72 | 26.37 | 156.19 | 26.64 | 21.12 | 32.17 | 391.93 | 12.30 | 39.59 | 708.14 | 31.07 | 14.81 | 3.54 |
| AL22 | 10.74 | 4.34 | 17.14 | 627.51 | 40.60 | 16.40 | 19.35 | 12.63 | 26.07 | 191.60 | 26.56 | 20.91 | 32.21 | 435.92 | 12.47 | 39.46 | 731.88 | 31.61 | 22.74 | 3.40 |
| AL79 | 10.30 | 4.12 | 16.48 | 743.63 | 39.80 | 17.40 | 19.18 | 12.73 | 25.63 | 230.98 | 26.57 | 21.07 | 32.07 | 410.90 | 12.04 | 37.90 | 749.24 | 31.77 | 24.23 | 3.19 |
| AL21 | 10.45 | 5.32 | 15.59 | 757.26 | 30.20 | 18.00 | 19.27 | 13.65 | 24.88 | 220.04 | 26.34 | 21.55 | 31.13 | 443.60 | 10.32 | 34.34 | 740.00 | 30.07 | 25.82 | 2.94 |
| MS8 | 9.03 | 3.23 | 14.83 | 751.16 | 44.20 | 26.60 | 18.96 | 12.93 | 25.00 | 205.60 | 26.75 | 21.39 | 32.12 | 366.89 | 11.59 | 35.04 | 817.35 | 33.08 | 30.54 | 2.21 |
| GA1 | 8.28 | 2.45 | 14.11 | 646.62 | 51.20 | 28.20 | 17.52 | 10.89 | 24.16 | 146.76 | 25.24 | 19.61 | 30.86 | 333.53 | 11.99 | 36.93 | 773.44 | 32.48 | 24.77 | 0.95 |
| ALBG | 7.98 | 2.43 | 13.52 | 803.97 | 53.40 | 34.00 | 17.36 | 11.28 | 23.43 | 205.16 | 25.05 | 19.78 | 30.33 | 375.49 | 11.37 | 35.25 | 787.57 | 32.26 | 28.87 | 0.86 |
| TN3 | 7.67 | 2.60 | 12.73 | 675.94 | 44.80 | 34.40 | 18.46 | 13.02 | 23.91 | 193.33 | 26.59 | 21.58 | 31.59 | 274.90 | 10.60 | 31.77 | 871.38 | 33.37 | 34.51 | 0.84 |
| TN34 | 6.84 | 0.49 | 13.20 | 706.96 | 69.40 | 41.80 | 16.24 | 9.11 | 23.38 | 189.93 | 25.01 | 18.70 | 31.32 | 400.29 | 13.18 | 37.70 | 829.34 | 34.97 | 26.60 | 0.26 |
| AR2 | 5.37 | -0.46 | 11.21 | 408.07 | 75.20 | 55.20 | 16.01 | 10.29 | 21.73 | 277.09 | 24.88 | 19.47 | 30.29 | 304.75 | 11.55 | 32.66 | 896.39 | 35.37 | 39.55 | -1.45 |
| KY5 | 4.65 | -0.76 | 10.05 | 639.24 | 76.40 | 62.20 | 14.93 | 8.74 | 21.12 | 186.46 | 23.50 | 17.89 | 29.11 | 424.26 | 11.56 | 33.74 | 858.56 | 34.26 | 20.69 | -1.84 |
| KY1 | 3.97 | -1.19 | 9.12 | 518.98 | 80.20 | 68.20 | 14.69 | 8.90 | 20.48 | 157.17 | 23.58 | 18.15 | 29.01 | 387.16 | 11.06 | 31.68 | 895.37 | 34.90 | 20.17 | -2.66 |
| MO2 | 2.64 | -2.75 | 8.03 | 331.60 | 87.60 | 73.60 | 15.09 | 9.32 | 20.85 | 178.42 | 24.70 | 19.08 | 30.33 | 315.03 | 11.44 | 30.22 | 1006.40 | 37.85 | 35.34 | -3.28 |
| OH1 | 3.18 | -1.77 | 8.13 | 441.33 | 86.00 | 74.80 | 14.25 | 8.44 | 20.06 | 179.55 | 23.43 | 18.02 | 28.84 | 384.76 | 10.78 | 30.80 | 922.27 | 35.00 | 22.41 | -3.29 |
| KS60 | 2.07 | -3.97 | 8.11 | 194.38 | 103.80 | 80.00 | 14.91 | 8.93 | 20.89 | 181.55 | 25.09 | 19.55 | 30.62 | 336.89 | 12.02 | 30.58 | 1042.56 | 39.32 | 61.91 | -3.79 |
| IN5 | 2.49 | -2.47 | 7.45 | 494.78 | 93.00 | 81.00 | 13.94 | 7.99 | 19.89 | 158.25 | 23.34 | 17.76 | 28.92 | 380.28 | 11.05 | 30.59 | 956.30 | 36.13 | 21.21 | -3.76 |
| OH119 | 1.74 | -3.10 | 6.57 | 402.92 | 96.60 | 88.20 | 13.30 | 7.63 | 18.96 | 160.35 | 22.94 | 17.49 | 28.40 | 323.92 | 10.73 | 29.88 | 965.83 | 35.90 | 22.42 | -4.67 |
| IN4 | 0.45 | -4.09 | 5.00 | 315.31 | 102.20 | 96.60 | 12.86 | 7.17 | 18.55 | 150.08 | 22.73 | 17.07 | 28.38 | 261.93 | 10.73 | 28.96 | 1019.50 | 37.06 | 27.80 | -5.78 |
| IA19 | -1.56 | -6.11 | 2.99 | 245.64 | 113.60 | 106.40 | 12.02 | 6.58 | 17.46 | 163.73 | 22.87 | 17.68 | 28.05 | 271.78 | 10.19 | 25.56 | 1113.19 | 39.86 | 40.63 | -7.27 |
| *Common gardens* | | | | | | | | | | | | | | | | | | | | |
| CG-RearS | 13.63 | 8.48 | 20.51 | 698.13 | 4.00 | 2.00 | 19.23 | 13.91 | 25.32 | 180.33 | 25.68 | 21.87 | 30.61 | 464.22 | 11.04 | 38.38 | 588.13 | 28.76 | 45.95 | 4.35 |
| CG-RearN | 8.50 | 0.68 | 16.07 | 707.66 | 79.00 | 27.00 | 16.68 | 10.77 | 24.12 | 285.21 | 22.94 | 18.76 | 28.67 | 493.39 | 12.71 | 39.25 | 711.61 | 32.37 | 57.53 | 0.72 |
| CG-Center | 6.46 | 0.91 | 14.15 | 514.40 | 69.00 | 53.00 | 16.42 | 10.33 | 22.94 | 122.99 | 22.14 | 18.32 | 26.77 | 392.85 | 11.95 | 37.67 | 716.13 | 31.72 | 47.85 | -1.16 |

Variable calculation detailed in Perrier et al. 2026.

**Table S3C: Summary of the principal component analysis (PCA) on seasonal and bioclimatic variables – Q1**

| | | PC1 | PC2 | PC3 | PC4 |
|---|---|---|---|---|---|
| Variance explained (%) | | 76.65 | 8.81 | 5.86 | 4.59 |
| Cumulative variance (%) | | 76.65 | 85.46 | 91.32 | 95.90 |
| *Loadings* | | | | | |
| Winter | Tmean | **0.26** | -0.02 | 0.00 | 0.02 |
| | Tmin | **0.26** | -0.05 | -0.06 | 0.09 |
| | Tmax | **0.26** | 0.00 | 0.05 | -0.04 |
| | PPTsum | 0.16 | -0.22 | **-0.51** | **-0.31** |
| | DayFrost | **-0.26** | 0.04 | 0.10 | -0.13 |
| | DayVerna | **-0.26** | 0.00 | 0.04 | 0.03 |
| Spring | Tmean | **0.26** | 0.08 | 0.00 | 0.12 |
| | Tmin | 0.25 | 0.11 | -0.10 | 0.24 |
| | Tmax | **0.26** | 0.05 | 0.08 | 0.02 |
| | PPTsum | 0.08 | **0.40** | **-0.54** | **-0.49** |
| Summer | Tmean | 0.25 | 0.23 | 0.02 | 0.13 |
| | Tmin | 0.24 | 0.24 | -0.08 | 0.25 |
| | Tmax | 0.25 | 0.21 | 0.12 | 0.01 |
| | PPTsum | 0.20 | -0.21 | 0.24 | 0.02 |
| Bioclim. | Bio 2 (Mean diurnal range) | 0.15 | 0.08 | **0.50** | **-0.64** |
| | Bio 3 (Isothermality) | 0.25 | -0.08 | 0.23 | -0.22 |
| | Bio 4 (Temperature seasonality) | -0.25 | 0.16 | -0.01 | 0.05 |
| | Bio 7 (Temperature annual range) | -0.25 | 0.20 | 0.11 | -0.11 |
| | Bio 15 (Precipitation seasonality | 0.08 | **0.70** | 0.14 | 0.10 |

Data only shown for principal components (PC) 1 to 4, as the cumulative contribution of the remaining PCs explained less than 5% of the variance. Loadings highlighted in bold indicate the main contributing variables to a given PC axis (>0.25).

**Table S4 Population average performance for traits and lifetime fitness for populations and common gardens (CG) – Q1**

| | Survival over winter | | | Bolting | | | Survival until reproduction | | | Flowering | | | Reproductive output | | | Lifetime fitness | | |
|---|---|---|---|---|---|---|---|---|---|---|---|---|---|---|---|---|---|---|
| | CG-Center | CG-RearN | CG-RearS | CG-Center | CG-RearN | CG-RearS | CG-Center | CG-RearN | CG-RearS | CG-Center | CG-RearN | CG-RearS | CG-Center | CG-RearN | CG-RearS | CG-Center | CG-RearN | CG-RearS |
| FL83 | 0.77 | 0.67 | 0.60 | 1.00 | 1.00 | 1.00 | 0.58 | 1.00 | 0.87 | 0.89 | 1.00 | 1.00 | 1.52 | 1.46 | 2.02 | 0.89 | 1.20 | 1.43 |
| FL81 | 0.70 | 0.63 | 0.30 | 1.00 | 1.00 | 1.00 | 0.46 | 1.00 | 0.83 | 0.75 | 0.97 | 0.90 | 1.01 | 1.08 | 1.63 | 0.32 | 0.83 | 0.66 |
| AL2 | 0.75 | 0.87 | 0.57 | 1.00 | 1.00 | 1.00 | 0.63 | 0.90 | 0.93 | 0.91 | 1.00 | 1.00 | 1.53 | 1.47 | 1.91 | 0.76 | 1.32 | 1.55 |
| MS6 | 0.93 | 0.73 | 0.70 | 1.00 | 1.00 | 0.93 | 0.73 | 0.93 | 0.97 | 1.00 | 1.00 | 1.00 | 1.61 | 1.57 | 2.08 | 1.35 | 1.42 | 1.80 |
| AL23 | 1.00 | 0.83 | 0.87 | 1.00 | 1.00 | 1.00 | 0.77 | 0.97 | 0.87 | 1.00 | 1.00 | 1.00 | 1.24 | 1.42 | 1.94 | 1.02 | 1.28 | 1.75 |
| AL22 | 0.87 | 0.63 | 0.70 | 1.00 | 1.00 | 1.00 | 0.69 | 0.93 | 0.83 | 1.00 | 1.00 | 1.00 | 1.51 | 1.36 | 2.00 | 1.03 | 1.04 | 1.60 |
| AL79 | 0.90 | 0.73 | 0.58 | 1.00 | 1.00 | 1.00 | 0.69 | 0.97 | 0.90 | 1.00 | 1.00 | 1.00 | 1.35 | 1.42 | 1.90 | 1.00 | 1.27 | 1.36 |
| AL21 | 0.90 | 0.73 | 0.53 | 1.00 | 0.97 | 0.42 | 0.65 | 0.97 | 0.90 | 0.65 | 1.00 | 0.93 | 1.42 | 1.54 | 1.90 | 0.78 | 1.42 | 0.59 |
| MS8 | 0.93 | 0.60 | 0.77 | 1.00 | 1.00 | 0.35 | 0.73 | 0.93 | 0.86 | 0.95 | 1.00 | 1.00 | 1.54 | 1.45 | 1.97 | 1.23 | 1.12 | 0.83 |
| GA1 | 0.87 | 0.79 | 0.75 | 1.00 | 1.00 | 0.14 | 0.39 | 1.00 | 0.87 | 1.00 | 1.00 | 0.93 | 1.36 | 1.62 | 1.94 | 0.56 | 1.46 | 0.31 |
| ALBG | 0.88 | 0.67 | 0.80 | 1.00 | 1.00 | 0.18 | 0.65 | 1.00 | 0.83 | 1.00 | 1.00 | 1.00 | 1.34 | 1.41 | 1.76 | 1.08 | 1.13 | 0.38 |
| TN3 | 0.93 | 0.83 | 0.90 | 1.00 | 1.00 | 0.27 | 0.85 | 1.00 | 0.93 | 1.00 | 1.00 | 1.00 | 1.53 | 1.47 | 2.00 | 1.37 | 1.41 | 0.81 |
| TN34 | 0.90 | 0.57 | 0.87 | 1.00 | 1.00 | 0.33 | 0.77 | 1.00 | 1.00 | 1.00 | 1.00 | 1.00 | 1.60 | 1.45 | 1.86 | 1.27 | 1.05 | 0.84 |
| AR2 | 0.97 | 0.63 | 0.70 | 1.00 | 1.00 | 0.31 | 0.69 | 1.00 | 0.87 | 1.00 | 1.00 | 0.93 | 1.52 | 1.39 | 1.90 | 1.24 | 1.00 | 0.50 |
| KY5 | 0.93 | 0.77 | 0.77 | 1.00 | 1.00 | 0.07 | 0.64 | 1.00 | 0.83 | 0.96 | 1.00 | 1.00 | 1.41 | 1.44 | 1.73 | 0.99 | 1.29 | 0.12 |
| KY1 | 0.92 | 0.79 | 0.67 | 1.00 | 1.00 | 0.00 | 0.59 | 0.93 | 0.63 | 1.00 | 0.97 | 0.86 | 1.49 | 1.42 | 1.88 | 1.06 | 1.29 | 0.00 |
| MO2 | 0.93 | 0.57 | 0.62 | 1.00 | 1.00 | 0.00 | 0.79 | 0.96 | 0.87 | 1.00 | 1.00 | 1.00 | 1.39 | 1.38 | 1.72 | 1.18 | 0.92 | 0.00 |
| OH1 | 0.87 | 0.83 | 0.97 | 1.00 | 1.00 | 0.00 | 0.88 | 1.00 | 0.90 | 1.00 | 1.00 | 0.93 | 1.66 | 1.50 | 1.92 | 1.53 | 1.37 | 0.00 |
| KS60 | 0.93 | 0.63 | 0.63 | 1.00 | 1.00 | 0.08 | 0.82 | 0.97 | 0.93 | 1.00 | 1.00 | 1.00 | 1.52 | 1.37 | 1.81 | 1.27 | 1.11 | 0.11 |
| IN5 | 1.00 | 0.67 | 0.89 | 1.00 | 1.00 | 0.00 | 0.73 | 0.93 | 0.90 | 0.86 | 1.00 | 0.96 | 1.72 | 1.45 | 1.87 | 1.30 | 1.13 | 0.00 |
| OH119 | 0.90 | 0.73 | 0.77 | 1.00 | 1.00 | 0.21 | 0.81 | 1.00 | 0.77 | 0.92 | 1.00 | 0.89 | 1.66 | 1.49 | 1.88 | 1.40 | 1.31 | 0.48 |
| IN4 | 0.97 | 0.80 | 0.93 | 1.00 | 1.00 | 0.07 | 0.92 | 0.93 | 0.90 | 1.00 | 1.00 | 0.89 | 1.46 | 1.45 | 1.88 | 1.42 | 1.36 | 0.11 |
| IA19 | 0.89 | 0.60 | 0.75 | 1.00 | 1.00 | 0.00 | 0.73 | 0.97 | 0.80 | 1.00 | 1.00 | 0.75 | 1.52 | 1.20 | 1.51 | 1.14 | 0.88 | 0.00 |

Seed-family level reproductive output and lifetime fitness were log10-transformed before calculating population averages.

**Table S5A: Correlation among life stages in the center garden – Q1**

| Life stages | Survival until reproduction | Flowering | Reproductive output |
|---|---|---|---|
| Survival over winter | 0.56 ** | 0.40 (*) | 0.35 |
| Survival until reproduction | - | 0.35 (*) | 0.50 * |
| Flowering | - | - | 0.25 |

Bolting was excluded in this garden because all plants bolted. Test statistics represent the Pearson's product–moment correlations between each trait pair. (*) P < 0.1; * P < 0.05; ** P < 0.01.

**Table S5B: Correlation among life stages in the northern rear-edge garden – Q1**

| Life stages | Bolting | Survival until reproduction | Flowering | Reproductive output |
|---|---|---|---|---|
| Survival over winter | -0.06 | -0.13 | 0.00 | 0.47 * |
| Bolting | - | 0.02 | -0.07 | -0.23 |
| Survival until reproduction | - | - | 0.03 | -0.08 |
| Flowering | - | - | - | 0.50 * |

Test statistics represent the Pearson's product–moment correlations between each trait pair. * P < 0.05.

**Table S5C: Correlation among life stages in the southern rear-edge garden – Q1**

| Life stages | Bolting | Survival until reproduction | Flowering | Reproductive output |
|---|---|---|---|---|
| Survival over winter | -0.49 * | 0.16 | 0.03 | 0.24 |
| Bolting | - | 0.23 | 0.38 (*) | 0.38 (*) |
| Survival until reproduction | - | - | 0.55 ** | 0.29 |
| Flowering | - | - | - | 0.54 ** |

Test statistics represent the Pearson's product–moment correlations between each trait pair. (*) P < 0.1; * P < 0.05; ** P < 0.01.

**Table S6: Variance inflation factors for life stages in each common garden (CG) – Q1**

| Life stages | CG-Center | CG-RearN | CG-RearS |
|---|---|---|---|
| Survival over winter | 1.57 | 1.44 | 1.94 |
| Bolting | - | 1.07 | 2.15 |
| Survival until reproduction | 1.76 | 1.02 | 1.53 |
| Flowering | 1.23 | 1.50 | 1.94 |
| Reproductive output | 1.35 | 2.02 | 1.93 |

Variance inflation factors were calculated for models testing for variation in fitness in linear models including all life stages as fixed effects. Bolting was excluded from the model in the center garden because all plants bolted.

**Table S7A: Description of variables calculated for question 2.**

| Variables | Description |
|---|---|
| Bolting | Individual-level scores of bolting (binary, success = 1) assessed in early and late spring in the four common gardens yielded estimates of bolting under eight distinct vernalization lengths for each population. For the 12 southernmost populations, bolting was also assessed under "no-vernalization" conditions in the greenhouse. |
| Bolting lag | The average number of days it takes a population to bolt after vernalization, measured under greenhouse conditions. |
| Vernalization length | Number of days with minimum air temperature < 4 °C during the period between the day of transplant and (day of visit - bolting lag) calculated for population for each of the early and late spring visits to the four common gardens. Set to zero for the no-vernalization treatment. |
| Vernalization requirement | Number of days of vernalization at which half the individuals of a population are predicted to have bolted, derived from population-level reaction norms of bolting vs vernalization length (Fig. 2A, Fig. S4, set to zero if negative). |
| Vernalization sensitivity | Slope of the bolting–vernalization length reaction norm on the logit scale. |

Variables were calculated at the level of the population with the exception of bolting.

**Table S7B: Description of variables calculated for question 3.**

| Variables | Description |
|---|---|
| Contemporary vernalization length | Number of days with minimum air temperature < 4 °C each year at a population's location of origin, averaged over 2018-2022.* |
| Future vernalization length | Number of days annually with predicted minimum air temperature < 4 °C for 2081-2100.** |
| Contemporary bolting probability | Bolting probability calculated by projecting *Contemporary vernalization length* on reaction norms of bolting vs vernalization length (Fig. 2A, Fig. S4). |
| Future bolting probability | Bolting probability calculated by projecting *Future vernalization length* on reaction norms of bolting vs vernalization length. |
| Bolting loss | (*Future bolting probability* - *Contemporary bolting probability*) / *Contemporary bolting probability*. |
| Expected change in vernalization length | *Contemporary vernalization length* - *Future vernalization length*. |
| Future vernalization length with fixed change | *Contemporary vernalization length* – absolute maximum *Expected change in vernalization length* across populations. |
| Future bolting probability with fixed change | *Future bolting probability* recalculated using *Future vernalization length with fixed change.* |
| Bolting loss with fixed change | *Bolting loss* recalculated using *Contemporary bolting probability* and *Future bolting probability with fixed change.* |
| Vernalization buffer length | *Contemporary vernalization length* - *Vernalization requirement* (Table S7A). |
| Contemporary vernalization length with fixed buffer | *Vernalization requirement* + minimum *Vernalization buffer length* across populations. |
| Future vernalization length with fixed buffer | *Contemporary vernalization length with fixed buffer* - *Expected change in vernalization length*. |
| Contemporary bolting probability with fixed buffer | *Contemporary bolting probability* recalculated using *Contemporary vernalization length with fixed buffer.* |
| Future bolting probability with fixed buffer | *Future bolting probability* recalculated using *Future vernalization length with fixed buffer.* |
| Bolting loss with fixed buffer | *Bolting loss* recalculated using *Contemporary bolting probability with fixed buffer* and *Future bolting probability with fixed buffer.* |

Variables were calculated for each population. "Future" variables were also calculated for each climate change scenario based on four Shared Socioeconomic Pathways (SSP). (*) Contemporary vernalization length was obtained from PRISM (https://prism.oregonstate.edu, accessed 26/01/2026). (**) Future vernalization length was estimated by converting projection of minimum temperature of the coldest month (BIO6) under future climates (2081-2100) obtained from Worldclim 2.1 (Fick & Hijmans, 2017) based on the relationship between Contemporary vernalization length and contemporary BIO6 (see Fig. S5).

**Table S8A: Population-average Bolting estimated for each population and vernalization length in the common gardens (CG) and in the greenhouse – Q2**

| | Bolting after short vernalization | | | | Bolting after long vernalization | | | | Bolting after no vernalization (greenhouse) |
|---|---|---|---|---|---|---|---|---|---|
| | CG-Center | CG-Int | CG-RearN | CG-RearS | CG-Center | CG-Int | CG-RearN | CG-RearS | |
| FL83 | 0.93 | 0.79 | 0.73 | 0.82 | 1.00 | 1.00 | 1.00 | 1.00 | 0.13 |
| FL81 | 0.57 | 0.23 | 0.20 | 0.70 | 1.00 | 1.00 | 1.00 | 1.00 | 0.89 |
| AL2 | 1.00 | 0.95 | 0.57 | 0.77 | 1.00 | 1.00 | 1.00 | 1.00 | 0.13 |
| MS6 | 0.97 | 0.86 | 0.60 | 0.83 | 1.00 | 1.00 | 1.00 | 0.93 | 0.00 |
| AL23 | 0.97 | 0.92 | 0.96 | 0.93 | 1.00 | 1.00 | 1.00 | 1.00 | 0.17 |
| AL22 | 0.77 | 0.79 | 0.77 | 0.92 | 1.00 | 1.00 | 1.00 | 1.00 | 0.33 |
| AL79 | 0.97 | 0.92 | 0.83 | 0.73 | 1.00 | 1.00 | 1.00 | 1.00 | 0.07 |
| AL21 | 0.90 | 0.82 | 0.50 | 0.12 | 1.00 | 1.00 | 0.97 | 0.42 | 0.03 |
| MS8 | 0.90 | 0.77 | 0.47 | 0.00 | 1.00 | 1.00 | 1.00 | 0.35 | 0.00 |
| GA1 | 0.79 | 0.85 | 0.08 | 0.00 | 1.00 | 1.00 | 1.00 | 0.14 | 0.03 |
| ALBG | 0.92 | 0.79 | 0.39 | 0.00 | 1.00 | 1.00 | 1.00 | 0.18 | 0.00 |
| TN3 | 0.60 | 0.63 | 0.23 | 0.00 | 1.00 | 1.00 | 1.00 | 0.27 | 0.00 |
| TN34 | 0.80 | 0.60 | 0.33 | 0.00 | 1.00 | 1.00 | 1.00 | 0.33 | - |
| AR2 | 0.57 | 0.68 | 0.03 | 0.00 | 1.00 | 1.00 | 1.00 | 0.31 | - |
| KY5 | 0.03 | 0.30 | 0.00 | 0.00 | 1.00 | 1.00 | 1.00 | 0.07 | - |
| KY1 | 0.12 | 0.23 | 0.00 | 0.00 | 1.00 | 1.00 | 1.00 | 0.00 | - |
| MO2 | 0.14 | 0.57 | 0.00 | 0.00 | 1.00 | 1.00 | 1.00 | 0.00 | - |
| OH1 | 0.73 | 0.61 | 0.13 | 0.00 | 1.00 | 1.00 | 1.00 | 0.00 | - |
| KS60 | 0.07 | 0.23 | 0.03 | 0.00 | 1.00 | 1.00 | 1.00 | 0.08 | - |
| IN5 | 0.27 | 0.54 | 0.07 | 0.00 | 1.00 | 1.00 | 1.00 | 0.00 | - |
| OH119 | 0.13 | 0.42 | 0.00 | 0.00 | 1.00 | 1.00 | 1.00 | 0.21 | - |
| IN4 | 0.27 | 0.77 | 0.07 | 0.00 | 1.00 | 1.00 | 1.00 | 0.07 | - |
| IA19 | 0.04 | 0.25 | 0.00 | 0.00 | 1.00 | 1.00 | 1.00 | 0.00 | - |

Bolting after short vernalization was quantified in early spring and after long vernalization quantified in late spring (Table S2B).

**Table S8B: Bolting lag and Vernalization length for each bolting measure after accounting for bolting lag – Q2**

| | Bolting lag (days) | Vernalization length in early spring (days) | | | | Vernalization length in late spring (days) | | | |
|---|---|---|---|---|---|---|---|---|---|
| | | CG-Center | CG-Int | CG-RearN | CG-RearS | CG-Center | CG-Int | CG-RearN | CG-RearS |
| FL83 | 29 | 92 | 87 | 81 | 34 | 116 | 105 | 97 | 37 |
| AL2 | 25 | 95 | 90 | 82 | 35 | 116 | 105 | 97 | 37 |
| MS6 | 24 | 95 | 90 | 83 | 35 | 116 | 105 | 97 | 37 |
| AL23 | 21 | 95 | 90 | 85 | 35 | 116 | 105 | 97 | 37 |
| AL22 | 19 | 95 | 90 | 86 | 35 | 116 | 105 | 97 | 37 |
| AL79 | 23 | 95 | 90 | 84 | 35 | 116 | 105 | 97 | 37 |
| AL21 | 29 | 92 | 87 | 81 | 34 | 116 | 105 | 97 | 37 |
| MS8 | 25 | 95 | 90 | 82 | 35 | 116 | 105 | 97 | 37 |
| GA1 | 34 | 89 | 85 | 78 | 32 | 115 | 105 | 97 | 37 |
| ALBG | 30 | 92 | 87 | 80 | 34 | 116 | 105 | 97 | 37 |
| TN3 | 31 | 92 | 87 | 80 | 34 | 116 | 105 | 97 | 37 |
| TN34 | 29 | 92 | 87 | 81 | 34 | 116 | 105 | 97 | 37 |
| AR2 | 27 | 94 | 89 | 82 | 35 | 116 | 105 | 97 | 37 |
| KY5* | 30 | 92 | 87 | 80 | 34 | 116 | 105 | 97 | 37 |
| KY1* | 31 | 92 | 87 | 80 | 34 | 116 | 105 | 97 | 37 |
| MO2* | 31 | 92 | 87 | 80 | 34 | 116 | 105 | 97 | 37 |
| OH1* | 31 | 92 | 87 | 80 | 34 | 116 | 105 | 97 | 37 |
| KS60 | 28 | 93 | 88 | 82 | 35 | 116 | 105 | 97 | 37 |
| IN5 | 34 | 89 | 85 | 78 | 32 | 115 | 105 | 97 | 37 |
| OH119 | 33 | 90 | 85 | 79 | 33 | 116 | 105 | 97 | 37 |
| IN4 | 27 | 94 | 89 | 82 | 35 | 116 | 105 | 97 | 37 |
| IA19 | 37 | 87 | 82 | 75 | 30 | 114 | 103 | 97 | 37 |

Bolting lag calculated from Perrier et al. (2025). For populations not included in that study (*), bolting lag was imputed based on the relationship between bolting lag and latitude of origin of populations with data (see Fig S3).

**Table S9A: Vernalization requirement and sensitivity – Q2**

| | Vernalization requirement (days) | Vernalization sensitivity (slope) |
|---|---|---|
| FL83 | 22.75 | 0.04 |
| AL2 | 24.60 | 0.04 |
| MS6 | 28.46 | 0.04 |
| AL23 | 10.01 | 0.05 |
| AL22* | 0.00 | 0.02 |
| AL79 | 23.95 | 0.04 |
| AL21 | 61.42 | 0.06 |
| MS8 | 64.40 | 0.07 |
| GA1 | 75.52 | 0.09 |
| ALBG | 72.27 | 0.09 |
| TN3 | 76.99 | 0.07 |
| TN34 | 70.36 | 0.06 |
| AR2 | 82.94 | 0.08 |
| KY5 | 93.48 | 0.23 |
| KY1 | 93.60 | 0.34 |
| MO2 | 91.60 | 0.25 |
| OH1 | 86.62 | 0.23 |
| KS60 | 94.27 | 0.23 |
| IN5 | 88.60 | 0.23 |
| OH119 | 88.49 | 0.09 |
| IN4 | 89.00 | 0.13 |
| IA19 | 90.63 | 0.27 |

* AL22 had bolting >0.5 under zero days of vernalization, resulting in negative values of vernalization requirement, which was therefore set to 0.

**Table S9B: Model comparisons between linear and quadratic effects of latitude – Q2**

| Dependent variable | *N* | $AICc_{\text{linear}}$ | $AICc_{\text{quadratic}}$ |
|---|---|---|---|
| Vernalization requirement | 22 | 189.75 | **181.68** |
| Vernalization sensitivity | 22 | **-55.26** | -53.23 |

The best model (bold) was identified by the lowest AICc value.

**Table S10A: Contemporary and Future vernalization length and minimum temperature of the coldest month for each population – Q3**

| Pop. | Contemporary vernalization length (days) 2018-2022 | Minimum Temperature (°C) of the Coldest Month | | | | | Future vernalization length (days) 2081–2100 | | | |
|---|---|---|---|---|---|---|---|---|---|---|
| | | 1970-2020 | 2081–2100 | | | | | | | |
| | | | SSP126 | SSP245 | SSP370 | SSP585 | SSP126 | SSP245 | SSP370 | SSP585 |
| FL83 | 55.48 | 4.00 | 4.30 | 5.40 | 7.60 | 6.70 | 60.55 | 54.42 | 42.17 | 47.18 |
| AL2 | 79.86 | 2.30 | 2.90 | 3.90 | 6.00 | 5.10 | 68.34 | 62.78 | 51.08 | 56.09 |
| MS6 | 83.33 | 0.80 | 1.80 | 3.00 | 4.60 | 4.20 | 74.47 | 67.79 | 58.88 | 61.11 |
| AL23 | 77.33 | 1.40 | 2.80 | 3.80 | 5.90 | 5.00 | 68.90 | 63.33 | 51.64 | 56.65 |
| AL22 | 82.29 | 0.80 | 1.80 | 3.00 | 4.80 | 4.10 | 74.47 | 67.79 | 57.76 | 61.66 |
| AL79 | 86.62 | 0.00 | 1.10 | 2.50 | 4.10 | 3.70 | 78.37 | 70.57 | 61.66 | 63.89 |
| AL21 | 71.81 | 0.00 | 0.80 | 2.00 | 3.50 | 3.30 | 80.04 | 73.36 | 65.00 | 66.12 |
| MS8 | 93.71 | -2.00 | -0.30 | 1.60 | 2.80 | 2.90 | 86.16 | 75.58 | 68.90 | 68.34 |
| GA1 | 100.57 | -2.40 | -0.50 | 1.20 | 2.80 | 2.40 | 87.28 | 77.81 | 68.90 | 71.13 |
| ALBG | 101.05 | -2.40 | -0.80 | 0.90 | 2.10 | 2.20 | 88.95 | 79.48 | 72.80 | 72.24 |
| TN3 | 95.86 | -2.30 | -0.40 | 1.80 | 2.70 | 3.10 | 86.72 | 74.47 | 69.46 | 67.23 |
| TN34 | 114.05 | -4.40 | -2.20 | -0.20 | 1.10 | 1.40 | 96.74 | 85.61 | 78.37 | 76.70 |
| AR2 | 114.43 | -4.50 | -2.80 | -0.70 | 0.50 | 0.80 | 100.08 | 88.39 | 81.71 | 80.04 |
| KY5 | 120.57 | -5.50 | -3.10 | -0.90 | 0.60 | 1.00 | 101.75 | 89.50 | 81.15 | 78.92 |
| KY1 | 124.29 | -6.70 | -3.80 | -1.40 | 0.00 | 0.70 | 105.65 | 92.29 | 84.49 | 80.59 |
| MO2 | 130.10 | -8.20 | -5.10 | -2.40 | -1.30 | -0.60 | 112.89 | 97.86 | 91.73 | 87.83 |
| OH1 | 127.86 | -7.80 | -4.60 | -2.20 | -0.50 | 0.30 | 110.11 | 96.74 | 87.28 | 82.82 |
| KS60 | 136.52 | -8.60 | -5.00 | -2.70 | -1.60 | -0.70 | 112.33 | 99.53 | 93.40 | 88.39 |
| IN5 | 130.14 | -8.70 | -5.10 | -2.40 | -1.10 | -0.20 | 112.89 | 97.86 | 90.62 | 85.61 |
| OH119 | 133.43 | -9.10 | -5.20 | -3.10 | -0.70 | 0.00 | 113.45 | 101.75 | 88.39 | 84.49 |
| IN4 | 137.71 | -10.60 | -6.10 | -3.40 | -1.70 | -0.50 | 118.46 | 103.43 | 93.96 | 87.28 |
| IA19 | 142.95 | -12.50 | -7.60 | -4.10 | -2.80 | -1.50 | 126.81 | 107.32 | 100.08 | 92.85 |

For each populations' location of origin, minimum temperature of the coldest month (BIO6) estimated for contemporary climates (1970-2020) and projections of future climates (2081-2100) for four Shared Socioeconomic Pathways (SSP) obtained from Worldclim 2.1 (Fick & Hijmans, 2017). Contemporary vernalization length was obtained from PRISM (2018–2022, https://prism.oregonstate.edu, accessed 26/01/2026). Expected Future vernalization length was estimated from projections of future BIO6 based on the relationship between Contemporary vernalization length and contemporary BIO6 (see Fig. S5).

**Table S10B: Predicted Contemporary bolting probability, Future bolting probability, and Bolting loss – Q3**

| Pop. | Contemporary bolting probability | Future bolting probability | | | | Bolting loss | | | |
|---|---|---|---|---|---|---|---|---|---|
| | | SSP126 | SSP245 | SSP370 | SSP585 | SSP126 | SSP245 | SSP370 | SSP585 |
| FL83 | 0.79 | 0.82 | 0.79 | 0.69 | 0.73 | 0.04 | -0.01 | -0.13 | -0.08 |
| AL2 | 0.89 | 0.84 | 0.81 | 0.73 | 0.77 | -0.06 | -0.09 | -0.18 | -0.14 |
| MS6 | 0.90 | 0.86 | 0.83 | 0.77 | 0.79 | -0.04 | -0.08 | -0.14 | -0.13 |
| AL23 | 0.96 | 0.94 | 0.92 | 0.87 | 0.90 | -0.02 | -0.04 | -0.09 | -0.06 |
| AL22 | 0.89 | 0.87 | 0.85 | 0.82 | 0.83 | -0.02 | -0.04 | -0.08 | -0.07 |
| AL79 | 0.94 | 0.92 | 0.89 | 0.84 | 0.86 | -0.02 | -0.06 | -0.10 | -0.09 |
| AL21 | 0.65 | 0.76 | 0.68 | 0.56 | 0.57 | 0.16 | 0.03 | -0.15 | -0.13 |
| MS8 | 0.88 | 0.81 | 0.68 | 0.58 | 0.57 | -0.07 | -0.23 | -0.35 | -0.36 |
| GA1 | 0.90 | 0.74 | 0.55 | 0.36 | 0.41 | -0.18 | -0.39 | -0.60 | -0.55 |
| ALBG | 0.93 | 0.82 | 0.66 | 0.51 | 0.50 | -0.12 | -0.29 | -0.45 | -0.46 |
| TN3 | 0.80 | 0.67 | 0.45 | 0.37 | 0.33 | -0.16 | -0.43 | -0.54 | -0.59 |
| TN34 | 0.94 | 0.84 | 0.72 | 0.62 | 0.60 | -0.11 | -0.23 | -0.34 | -0.36 |
| AR2 | 0.92 | 0.79 | 0.60 | 0.48 | 0.45 | -0.14 | -0.34 | -0.48 | -0.51 |
| KY5 | 1.00 | 0.87 | 0.29 | 0.06 | 0.03 | -0.13 | -0.71 | -0.94 | -0.97 |
| KY1 | 1.00 | 0.98 | 0.39 | 0.04 | 0.01 | -0.02 | -0.61 | -0.96 | -0.99 |
| MO2 | 1.00 | 0.99 | 0.82 | 0.51 | 0.28 | -0.01 | -0.18 | -0.49 | -0.72 |
| OH1 | 1.00 | 1.00 | 0.91 | 0.54 | 0.29 | 0.00 | -0.09 | -0.46 | -0.71 |
| KS60 | 1.00 | 0.98 | 0.77 | 0.45 | 0.21 | -0.02 | -0.23 | -0.55 | -0.79 |
| IN5 | 1.00 | 1.00 | 0.89 | 0.61 | 0.34 | 0.00 | -0.11 | -0.39 | -0.66 |
| OH119 | 0.98 | 0.91 | 0.77 | 0.50 | 0.41 | -0.08 | -0.21 | -0.49 | -0.59 |
| IN4 | 1.00 | 0.98 | 0.87 | 0.66 | 0.44 | -0.02 | -0.13 | -0.34 | -0.56 |
| IA19 | 1.00 | 1.00 | 0.99 | 0.93 | 0.65 | 0.00 | -0.01 | -0.07 | -0.35 |

**Table S11A: Contemporary and Future vernalization length (days) estimates that account for spatial variation in climate – Q3**

| Pop. | Expected change in vernalization length * | | | | Future vernalization length with fixed change 2081–2100 | | | | Vernalization buffer length ** | Contemporary vernalization length with fixed buffer 2018–2022 | Future vernalization length with fixed buffer 2081–2100 | | | |
|---|---|---|---|---|---|---|---|---|---|---|---|---|---|---|
| | SSP126 | SSP245 | SSP370 | SSP585 | SSP126 | SSP245 | SSP370 | SSP585 | | | SSP126 | SSP245 | SSP370 | SSP585 |
| FL83 | -5.07 | 1.06 | 13.31 | 8.30 | 31.29 | 18.48 | 10.44 | 5.04 | 32.73 | 33.14 | 38.21 | 32.09 | 19.84 | 24.85 |
| AL2 | 11.52 | 17.08 | 28.78 | 23.77 | 55.67 | 42.86 | 34.82 | 29.42 | 55.26 | 34.99 | 23.48 | 17.91 | 6.22 | 11.23 |
| MS6 | 8.86 | 15.54 | 24.45 | 22.22 | 59.14 | 46.34 | 38.30 | 32.90 | 54.87 | 38.86 | 30.00 | 23.31 | 14.40 | 16.63 |
| AL23 | 8.43 | 14.00 | 25.69 | 20.68 | 53.14 | 40.34 | 32.30 | 26.90 | 67.33 | 20.40 | 11.97 | 6.40 | -5.29 | -0.28 |
| AL22 | 7.82 | 14.50 | 24.53 | 20.63 | 58.10 | 45.29 | 37.25 | 31.85 | 82.29 | 10.39 | 2.58 | -4.10 | -14.13 | -10.23 |
| AL79 | 8.25 | 16.05 | 24.96 | 22.73 | 62.43 | 49.62 | 41.58 | 36.18 | 62.67 | 34.34 | 26.09 | 18.29 | 9.38 | 11.61 |
| AL21 | -8.23 | -1.55 | 6.81 | 5.69 | 47.62 | 34.81 | 26.77 | 21.37 | **10.39** | 71.81 | 80.04 | 73.36 | 65.00 | 66.12 |
| MS8 | 7.55 | 18.13 | 24.81 | 25.37 | 69.53 | 56.72 | 48.68 | 43.28 | 29.31 | 74.80 | 67.25 | 56.67 | 49.98 | 49.43 |
| GA1 | 13.29 | 22.76 | 31.67 | 29.44 | 76.38 | 63.58 | 55.53 | 50.13 | 25.05 | 85.91 | 72.62 | 63.15 | 54.24 | 56.47 |
| ALBG | 12.10 | 21.57 | 28.25 | 28.81 | 76.86 | 64.05 | 56.01 | 50.61 | 28.78 | 82.66 | 70.56 | 61.10 | 54.41 | 53.86 |
| TN3 | 9.14 | 21.39 | 26.40 | 28.63 | 71.67 | 58.86 | 50.82 | 45.42 | 18.87 | 87.38 | 78.24 | 65.99 | 60.98 | 58.75 |
| TN34 | 17.31 | 28.44 | 35.68 | 37.35 | 89.86 | 77.05 | 69.01 | 63.61 | 43.69 | 80.75 | 63.45 | 52.31 | 45.07 | 43.40 |
| AR2 | 14.35 | 26.04 | 32.72 | 34.39 | 90.24 | 77.43 | 69.39 | 63.99 | 31.48 | 93.34 | 78.99 | 67.30 | 60.62 | 58.95 |
| KY5 | 18.82 | 31.07 | 39.42 | 41.65 | 96.38 | 83.58 | 75.53 | 70.13 | 27.09 | 103.87 | 85.06 | 72.81 | 64.45 | 62.23 |
| KY1 | 18.64 | 32.00 | 39.80 | 43.70 | 100.10 | 87.29 | 79.25 | 73.85 | 30.69 | 103.99 | 85.36 | 71.99 | 64.20 | 60.30 |
| MO2 | 17.21 | 32.24 | 38.37 | 42.27 | 105.91 | 93.10 | 85.06 | 79.66 | 38.50 | 101.99 | 84.79 | 69.75 | 63.63 | 59.73 |
| OH1 | 17.75 | 31.12 | 40.58 | 45.04 | 103.67 | 90.86 | 82.82 | 77.42 | 41.24 | 97.01 | 79.26 | 65.90 | 56.43 | 51.97 |
| KS60 | **24.19** | **36.99** | 43.12 | 48.13 | 112.33 | 99.53 | 91.49 | 86.09 | 42.26 | 104.66 | 80.47 | 67.66 | 61.54 | 56.53 |
| IN5 | 17.25 | 32.28 | 39.52 | 44.53 | 105.95 | 93.15 | 85.10 | 79.71 | 41.55 | 98.99 | 81.74 | 66.70 | 59.47 | 54.45 |
| OH119 | 19.98 | 31.68 | **45.04** | 48.94 | 109.24 | 96.43 | 88.39 | 82.99 | 44.94 | 98.88 | 78.90 | 67.21 | 53.84 | 49.94 |
| IN4 | 19.25 | 34.28 | 43.75 | **50.43** | 113.53 | 100.72 | 92.68 | 87.28 | 48.72 | 99.39 | 80.14 | 65.10 | 55.64 | 48.95 |
| IA19 | 16.14 | 35.63 | 42.87 | 50.10 | 118.76 | 105.96 | 97.91 | 92.52 | 52.32 | 101.03 | 84.89 | 65.40 | 58.16 | 50.92 |

Values in bold represent the populations with the highest Expected change in vernalization length (*) and the smallest Vernalization buffer length (**), used to calculate Future vernalization length with fixed change and fixed buffer.

**Table S11B: Predicted Future bolting probability and Bolting loss estimates that account for spatial variation in expected climate change – Q3**

| | Future bolting probability with fixed change | | | | Bolting loss with fixed change | | | |
|---|---|---|---|---|---|---|---|---|
| Pop. | SSP126 | SSP245 | SSP370 | SSP585 | SSP126 | SSP245 | SSP370 | SSP585 |
| FL83 | 0.59 | 0.46 | 0.38 | 0.33 | -0.26 | -0.42 | -0.53 | -0.59 |
| AL2 | 0.76 | 0.67 | 0.60 | 0.55 | -0.14 | -0.25 | -0.33 | -0.39 |
| MS6 | 0.77 | 0.67 | 0.60 | 0.54 | -0.14 | -0.25 | -0.34 | -0.39 |
| AL23 | 0.88 | 0.80 | 0.74 | 0.69 | -0.08 | -0.16 | -0.23 | -0.28 |
| AL22 | 0.82 | 0.77 | 0.74 | 0.71 | -0.08 | -0.13 | -0.17 | -0.20 |
| AL79 | 0.85 | 0.76 | 0.69 | 0.63 | -0.10 | -0.19 | -0.27 | -0.33 |
| AL21 | 0.30 | 0.16 | 0.11 | 0.08 | -0.54 | -0.75 | -0.84 | -0.88 |
| MS8 | 0.59 | 0.37 | 0.26 | 0.19 | -0.33 | -0.58 | -0.71 | -0.78 |
| GA1 | 0.52 | 0.26 | 0.15 | 0.10 | -0.42 | -0.71 | -0.83 | -0.89 |
| ALBG | 0.60 | 0.32 | 0.19 | 0.12 | -0.35 | -0.65 | -0.80 | -0.87 |
| TN3 | 0.40 | 0.21 | 0.13 | 0.09 | -0.49 | -0.74 | -0.84 | -0.89 |
| TN34 | 0.77 | 0.60 | 0.48 | 0.40 | -0.18 | -0.36 | -0.49 | -0.58 |
| AR2 | 0.63 | 0.40 | 0.26 | 0.19 | -0.31 | -0.57 | -0.71 | -0.79 |
| KY5 | 0.66 | 0.09 | 0.02 | 0.00 | -0.34 | -0.91 | -0.98 | -1.00 |
| KY1 | 0.90 | 0.11 | 0.01 | 0.00 | -0.10 | -0.89 | -0.99 | -1.00 |
| MO2 | 0.97 | 0.59 | 0.17 | 0.05 | -0.03 | -0.41 | -0.83 | -0.95 |
| OH1 | 0.98 | 0.73 | 0.29 | 0.11 | -0.02 | -0.27 | -0.71 | -0.89 |
| KS60 | 0.98 | 0.77 | 0.35 | 0.13 | -0.02 | -0.23 | -0.65 | -0.87 |
| IN5 | 0.98 | 0.74 | 0.31 | 0.12 | -0.02 | -0.26 | -0.69 | -0.88 |
| OH119 | 0.87 | 0.68 | 0.50 | 0.38 | -0.11 | -0.31 | -0.49 | -0.62 |
| IN4 | 0.96 | 0.83 | 0.62 | 0.44 | -0.03 | -0.17 | -0.38 | -0.56 |
| IA19 | 1.00 | 0.99 | 0.88 | 0.63 | 0.00 | -0.01 | -0.12 | -0.37 |

**Table S11C: Predicted Contemporary bolting probability, Future bolting probability, and Bolting loss estimates that account for spatial variation in the buffer to vernalization requirement – Q3**

| Pop. | Contemporary bolting probability with fixed buffer | Future bolting probability with fixed buffer | | | | Bolting loss with fixed buffer | | | |
|---|---|---|---|---|---|---|---|---|---|
| | | SSP126 | SSP245 | SSP370 | SSP585 | SSP126 | SSP245 | SSP370 | SSP585 |
| FL83 | 0.60 | 0.65 | 0.59 | 0.47 | 0.52 | 0.08 | -0.02 | -0.22 | -0.14 |
| AL2 | 0.60 | 0.49 | 0.44 | 0.33 | 0.38 | -0.18 | -0.27 | -0.44 | -0.37 |
| MS6 | 0.60 | 0.52 | 0.45 | 0.36 | 0.38 | -0.14 | -0.25 | -0.40 | -0.36 |
| AL23 | 0.62 | 0.52 | 0.46 | 0.33 | 0.38 | -0.15 | -0.26 | -0.47 | -0.38 |
| AL22 | 0.59 | 0.55 | 0.51 | 0.45 | 0.47 | -0.08 | -0.14 | -0.25 | -0.21 |
| AL79 | 0.61 | 0.52 | 0.44 | 0.34 | 0.37 | -0.15 | -0.29 | -0.44 | -0.41 |
| AL21 | 0.65 | 0.76 | 0.68 | 0.56 | 0.57 | 0.16 | 0.03 | -0.15 | -0.13 |
| MS8 | 0.67 | 0.55 | 0.37 | 0.27 | 0.27 | -0.18 | -0.44 | -0.59 | -0.60 |
| GA1 | 0.71 | 0.44 | 0.25 | 0.13 | 0.16 | -0.39 | -0.64 | -0.81 | -0.78 |
| ALBG | 0.72 | 0.46 | 0.27 | 0.17 | 0.16 | -0.36 | -0.63 | -0.77 | -0.78 |
| TN3 | 0.68 | 0.52 | 0.31 | 0.24 | 0.21 | -0.23 | -0.54 | -0.65 | -0.69 |
| TN34 | 0.65 | 0.40 | 0.25 | 0.17 | 0.16 | -0.40 | -0.62 | -0.73 | -0.76 |
| AR2 | 0.69 | 0.43 | 0.23 | 0.16 | 0.14 | -0.38 | -0.66 | -0.77 | -0.80 |
| KY5 | 0.92 | 0.13 | 0.01 | 0.00 | 0.00 | -0.86 | -0.99 | -1.00 | -1.00 |
| KY1 | 0.97 | 0.06 | 0.00 | 0.00 | 0.00 | -0.94 | -1.00 | -1.00 | -1.00 |
| MO2 | 0.93 | 0.16 | 0.00 | 0.00 | 0.00 | -0.83 | -0.99 | -1.00 | -1.00 |
| OH1 | 0.92 | 0.15 | 0.01 | 0.00 | 0.00 | -0.83 | -0.99 | -1.00 | -1.00 |
| KS60 | 0.92 | 0.04 | 0.00 | 0.00 | 0.00 | -0.96 | -1.00 | -1.00 | -1.00 |
| IN5 | 0.91 | 0.18 | 0.01 | 0.00 | 0.00 | -0.81 | -0.99 | -1.00 | -1.00 |
| OH119 | 0.72 | 0.29 | 0.12 | 0.04 | 0.03 | -0.60 | -0.83 | -0.95 | -0.96 |
| IN4 | 0.80 | 0.23 | 0.04 | 0.01 | 0.00 | -0.71 | -0.95 | -0.99 | -0.99 |
| IA19 | 0.94 | 0.17 | 0.00 | 0.00 | 0.00 | -0.82 | -1.00 | -1.00 | -1.00 |

**Table S12A. Estimates of the effect of ΔPC1 on performance – Q1**

| | CG-Center | | CG-RearN | | CG-RearS | |
|---|---|---|---|---|---|---|
| Life stages | $\beta$ ΔPC1 | $\beta$ $ΔPC1^2$ | $\beta$ ΔPC1 | $\beta$ $ΔPC1^2$ | $\beta$ ΔPC1 | $\beta$ $ΔPC1^2$ |
| Survival over winter | 0.006 | -0.003 | 0.000 | -0.001 | **0.057** ** | **-0.004** * |
| Bolting | 0.000 | 0.000 | 0.000 | 0.000 | **-0.185** *** | **0.009** *** |

Values depict the estimated slope coefficients ($\beta$) of the linear and square effect of ΔPC1 in each common garden (CG) for each life stage. *P*-values of the linear and square trends were estimated from the full model (Table 1; *t*-test at ΔPC1=0) and are indicated as (*) P < 0.1; * P < 0.05; ** P < 0.01; *** P < 0.001.

**Table S12B Comparison of the effect of ΔPC1 on performance between common gardens (CG) – Q1**

| | CG-Center vs CG-RearN | | CG-Center vs CG-RearS | | CG-RearN vs CG-RearS | |
|---|---|---|---|---|---|---|
| Life stages | $\beta$ CG * ΔPC1 | $\beta$ CG * $ΔPC1^2$ | $\beta$ CG * ΔPC1 | $\beta$ CG * $ΔPC1^2$ | $\beta$ CG * ΔPC1 | $\beta$ CG * $ΔPC1^2$ |
| Survival over winter | 0.006 | -0.002 | **-0.051** * | 0.001 | **-0.058** ** | 0.003 |
| Bolting | 0.000 | 0.000 | **0.185** *** | **-0.010** *** | **0.186** *** | **-0.010** *** |

Values depict the estimated pairwise contrast ($\beta$) of the linear and quadratic effect of ΔPC1 between common gardens (based on the garden - ΔPC1 interaction) obtained from Tukey's tests for each life stage. Estimates with *P*-values * < 0.05; ** *P*<0.01, *** *P*<0.001.

**Table S13: Variation in Bolting across Vernalization length - Q2**

| Fixed effects | $\beta$ | $X^2$ | | $R^2_m$ | $R^2_c$ |
|---|---|---|---|---|---|
| Vernalization length | 0.13 | 34.13 | *** | 0.30 | 0.58 |

Bolting was tested in a linear model with Vernalization length as continuous predictor. Test statistics include the slope of the effect of Vernalization length, the chi-squared value of the model, and the marginal ($R^2$m, fixed effect only) and conditional ($R^2$c, fixed and random effects) $R^2$. Random effects not shown. *** P < 0.001.

**Table S14: Variation in Vernalization requirement and sensitivity across latitude – Q2**

| Dependent variable | $\beta$ latitude | $\beta$ latitude$^2$ | $F$ | | $R^2$ |
|---|---|---|---|---|---|
| Vernalization requirement | 123.25 | -45.11 | 52.70 | *** | 0.83 |
| Vernalization sensitivity | 0.02 | - | 31.12 | *** | 0.59 |

Dependent variables were tested in separate linear models with latitude either as linear or quadratic continuous predictor (model selection in table S9B). Test statistics include the slope of the effect of latitude ($\beta$) estimated for the linear and the squared term (if applicable), the $F$ value and the $R^2$ of each model. *** P < 0.001.

**Table S15: Variation in Bolting loss with fixed change across latitude for each climate change scenario – Q3**

| | *β* latitude | *β* latitude$^2$ | *F* | | $R^2$ |
|---|---|---|---|---|---|
| SSP126 | 0.32 | 0.38 | 7.04 | ** | 0.37 |
| SSP245 | 0.11 | 0.79 | 7.58 | ** | 0.39 |
| SSP370 | -0.24 | 0.83 | 9.77 | ** | 0.46 |
| SSP585 | -0.47 | 0.70 | 10.12 | ** | 0.46 |

Bolting loss expected under future climates assuming a fixed change in vernalization length across the range (Table S7b) was estimated for climate change scenarios representing a range of future warming, and tested in separate linear models for each scenario, with latitude as continuous quadratic predictor. Test statistics include the slope of the effect of latitude (*β*) estimated for the linear and the squared term, the *F* value and the $R^2$ of each model. ** P < 0.01; *** P < 0.001.

**Table S16: Variation in Bolting loss with fixed buffer across latitude for each climate change scenario – Q3**

| | *β* latitude | *β* latitude$^2$ | *F* | | $R^2$ |
|---|---|---|---|---|---|
| SSP126 | -1.42 | 0.11 | 34.79 | *** | 0.76 |
| SSP245 | -1.51 | 0.25 | 61.46 | *** | 0.85 |
| SSP370 | -1.21 | 0.23 | 50.14 | *** | 0.82 |
| SSP585 | -1.32 | 0.31 | 66.31 | *** | 0.86 |

Bolting loss expected under future climates assuming a fixed vernalization buffer across the range (Table S7b) was estimated for climate change scenarios representing a range of future warming, and tested in separate linear models for each scenario, with latitude as continuous quadratic predictor. Test statistics include the slope of the effect of latitude (*β*) estimated for the linear and the squared term, the *F* value and the $R^2$ of each model. *** P < 0.001.

**Figures**

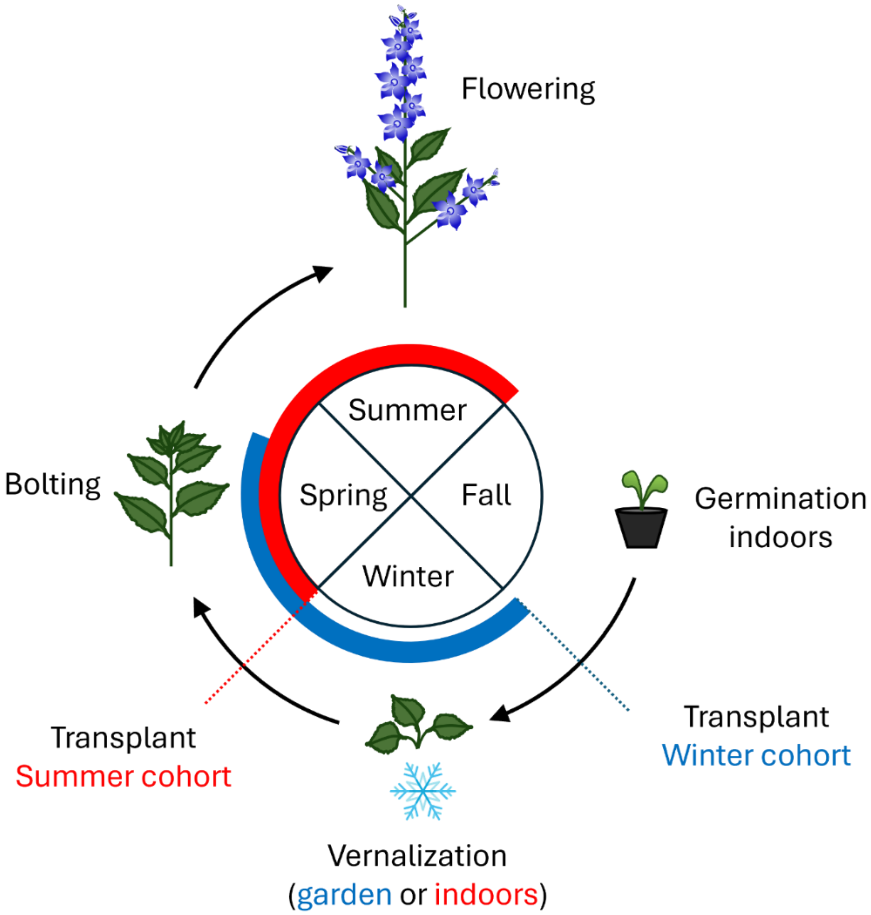

**Figure S1: Life cycle of *C. americana* split between winter and summer cohorts (adapted from Perrier et al. 2026).** The common garden experiment consisted of a summer cohort (red, spring and summer 2023) and a winter cohort (blue, fall 2023 – spring 2024). Plants were germinated indoors for both cohorts, then transplanted into each garden before winter for the winter cohort, and after a simulated winter indoors for the summer cohort. Together, the two cohorts allowed determining the effects of climate across most of the species' life cycle.

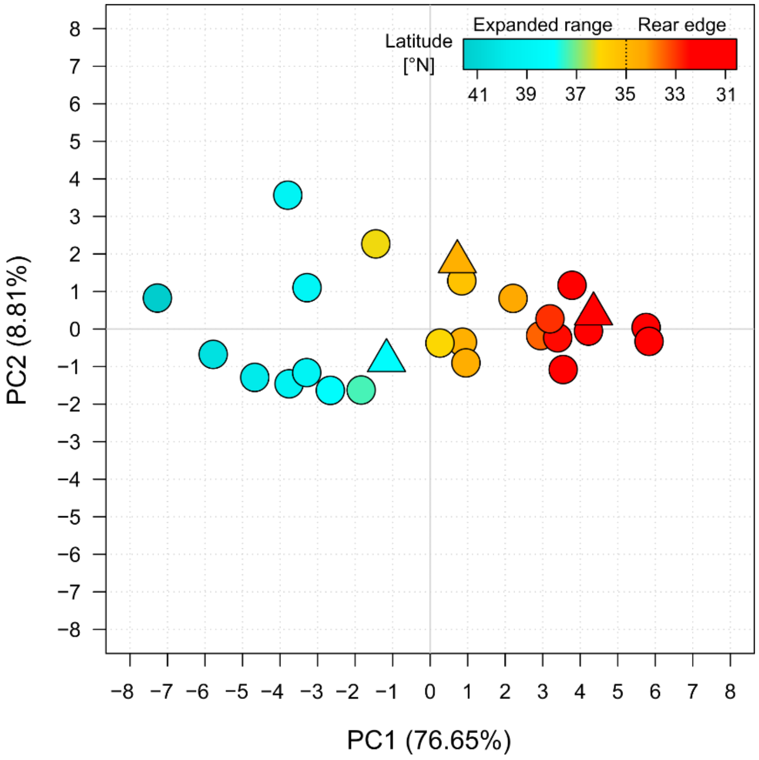

**Figure S2. Climate conditions in each common garden were generally comparable to average conditions in nearby populations**. Climate variation across the 23 populations (dots) included in the experiment is represented by their relative position along the first two axes of a principal component analysis based on 19 seasonal and bioclimatic variables estimated for each population's location of origin. Common gardens (triangles, cyan, CG-Center; yellow, CG-Int; red, CG-Rear) were projected onto the populations' PC space based on the same climate variables. Populations are colored by latitude.

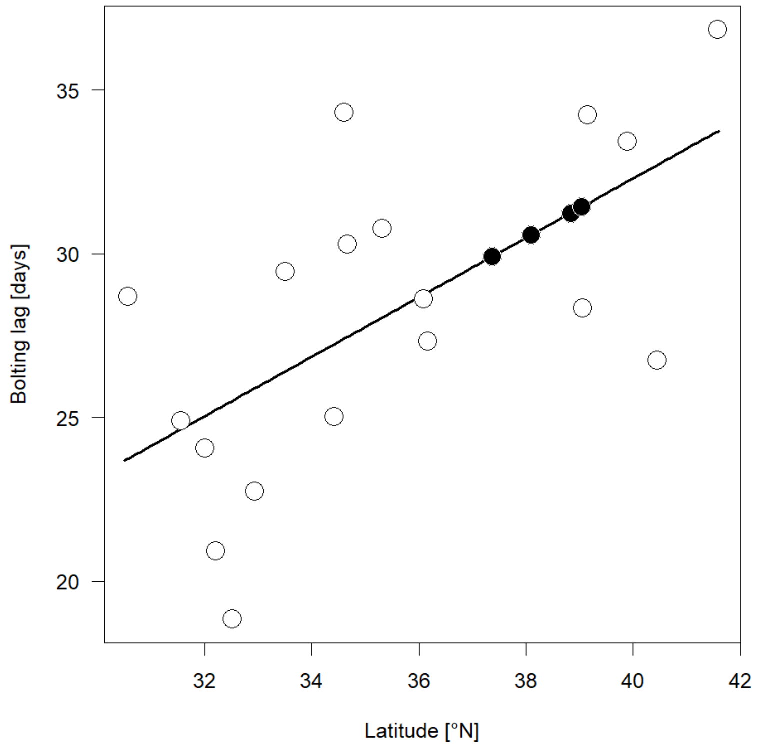

**Figure S3: Estimation of bolting lag.** Population-specific bolting lag (white dots) was estimated for each population as the average number of days from the end of vernalization to bolting from previously published data (Perrier et al. 2025). This data was not available for four populations and was imputed (black dots) based on the relationship between the measured bolting lag and populations' latitude of origin (black line, $R^2 = 0.36$, $P = 0.005$).

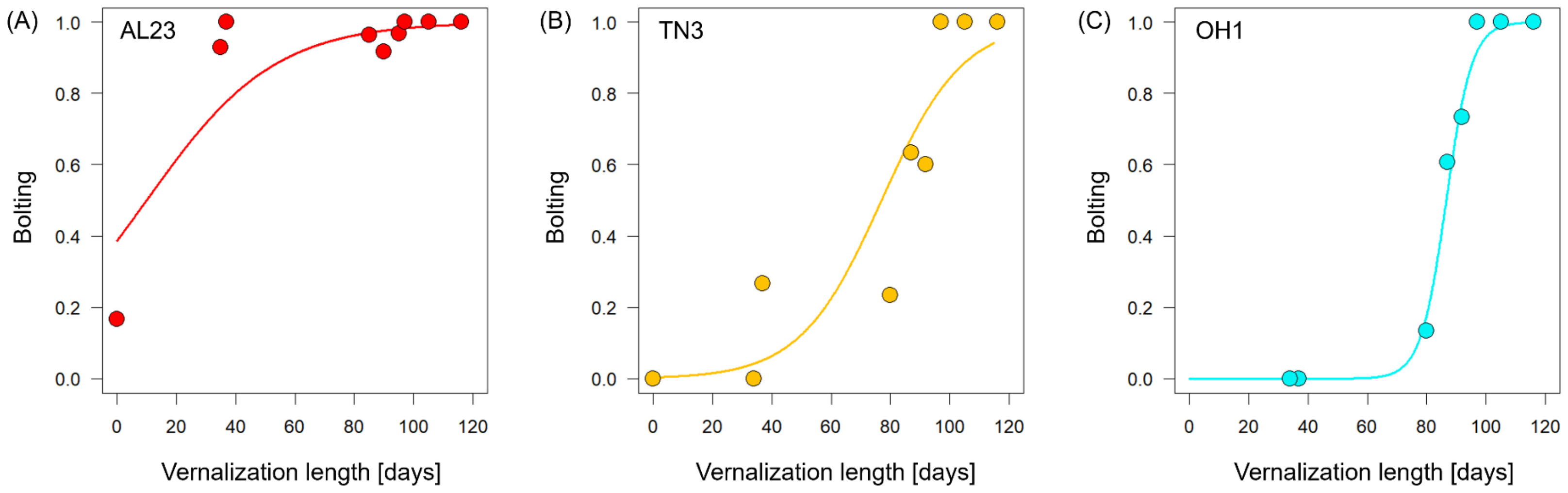

**Figure S4: Bolting-Vernalization length reaction norms for representative populations from (A) southern and (B) northern rear-edge and (C) central latitudes**. Lines represent population-specific model-predicted relationship between bolting and vernalization length, with population-average bolting in each vernalization length indicated as dots. Test statistics are reported in Table S12.

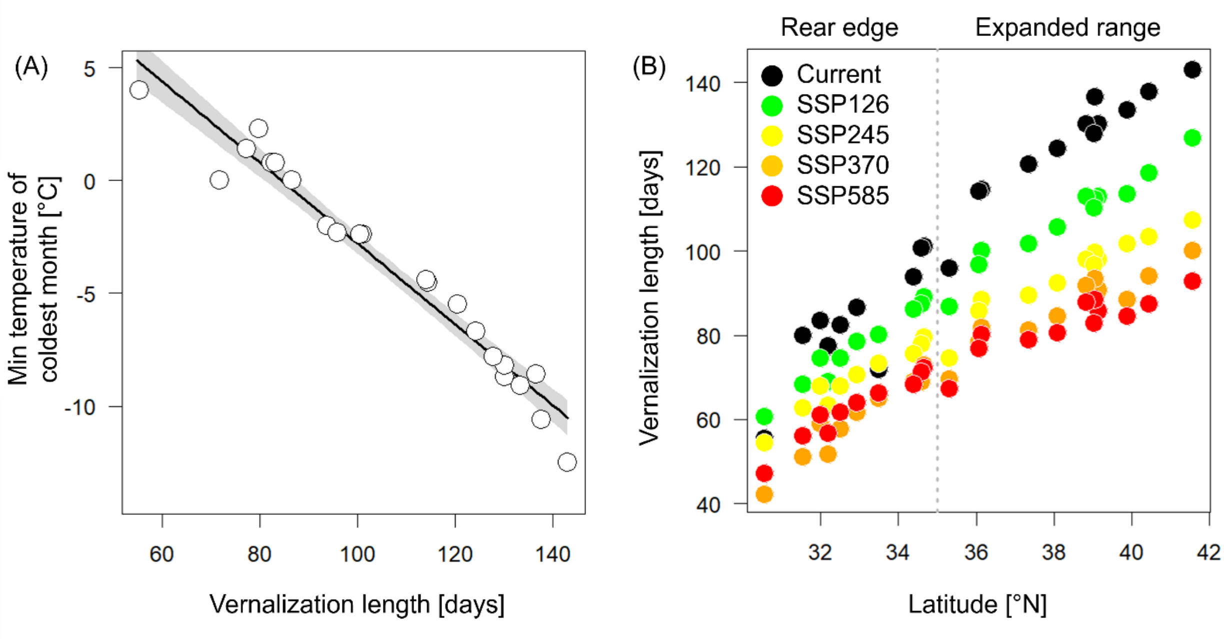

**Figure S5: Estimation of future vernalization length. (A)** The black line represents the relationship between Minimum Temperature of the Coldest Month (BIO6) from WorldClim 2.1 data (1970-2020, Fick & Hijmans, 2017) and vernalization length estimated from recent daily PRISM temperature data (2018–2022, https://prism.oregonstate.edu ) for the location of origin of each population (white dots), with the 95% CI indicated in gray shading ($R^2$ = 0.96). **(B)** Vernalization length across latitude for each population (dots) based on current PRISM data (black), and estimated vernalization length under future climates (2081–2100) from BIO6 estimated from WorldClim 2.1 for four Shared Socio-economic Pathways (green: SSP126, +1.8 °C; yellow: SSP245, +2.7 °C; orange: SSP370, +3.6 °C; red: SSP585, +4.4°C) converted based on the relationship in (A). The vertical dotted line represents the separation between the rear edge (<35°N) and the expanded range.

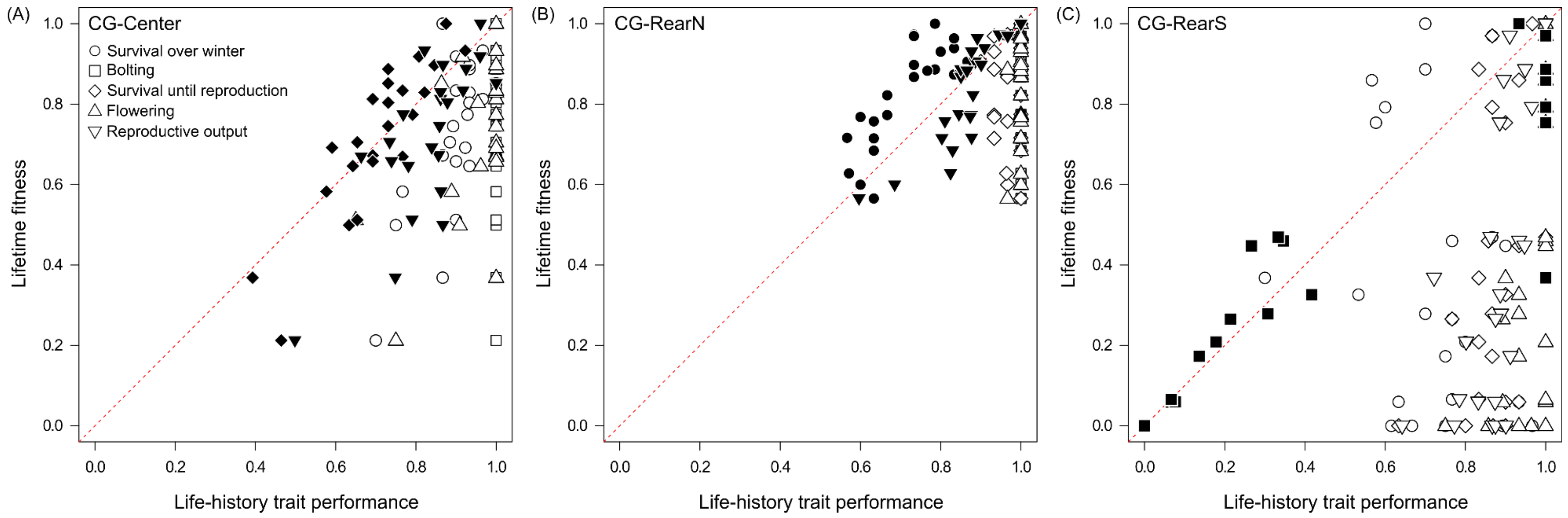

**Figure S6: Bolting contributes the most to lifetime fitness at the rear edge.** Variation in lifetime fitness of populations (individual symbols) raised in the center (A), northern rear-edge (B) and (C) southern rear-edge common garden relative to performance for each life- stage (distinguished by symbols). The traits contributing the most to lifetime fitness are indicated in black (Table 2). Red lines represent the 1:1 relationship between lifetime fitness and traits.

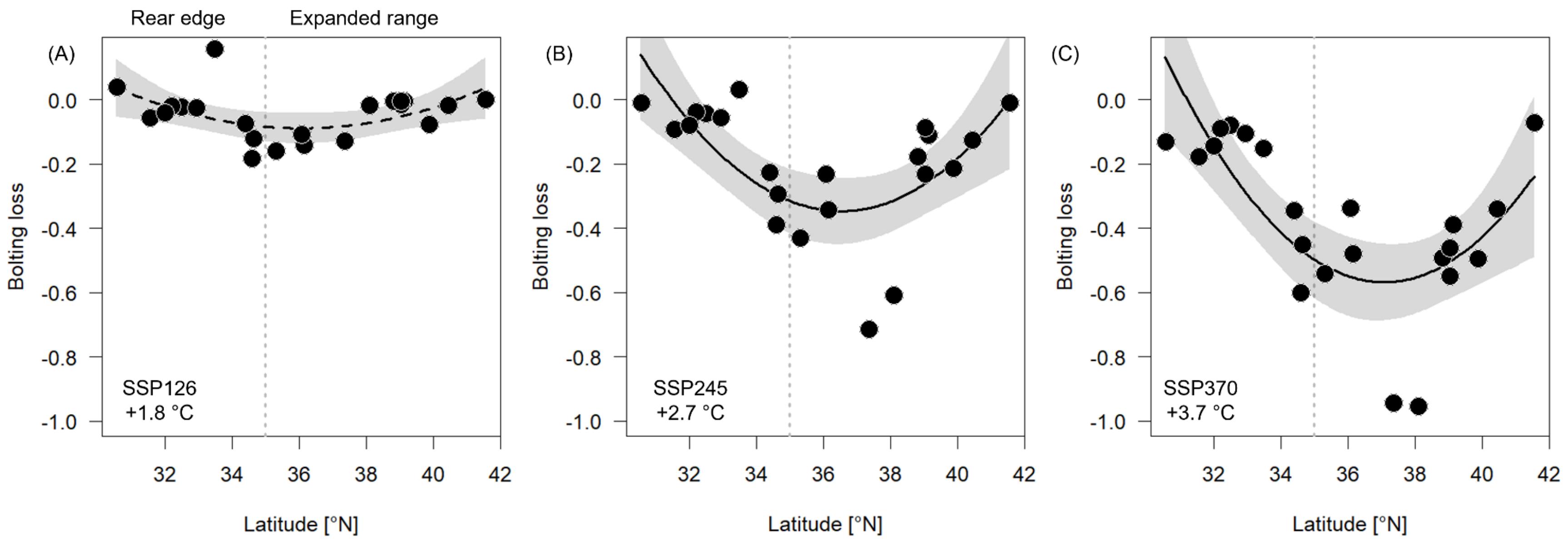

**Figure S7: Bolting loss under future climates (2081–2100)**. Bolting loss estimated for three Shared Socioeconomic Pathways for each population (dots). **(A)** SSP126, +1.8 °C, **(B)** SSP245, +2.7 °C, **(C)** SSP370, +3.6 °C. The vertical dotted line represents the separation between the rear edge (<35°N) and the expanded range. The solid line represents the significant relationship between each estimate of bolting loss and latitude (dashed if $P < 0.05$), with the 95% CI indicated in gray shading. Test statistics are reported in Table 3.

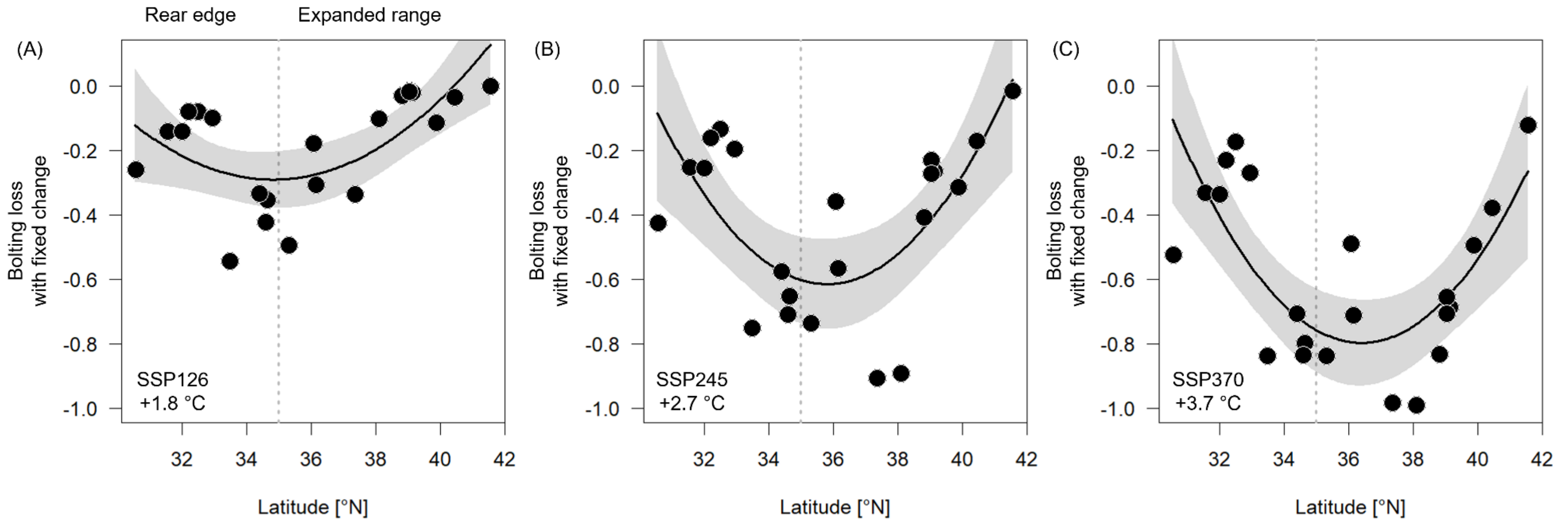

**Figure S8: Bolting loss with fixed change under future climates (2081–2100).** Bolting loss estimated for three Shared Socioeconomic Pathways for each population (dots), with expected change in vernalization held constant across the range. **(A)** SSP126, +1.8 °C, **(B)** SSP245, +2.7 °C, **(C)** SSP370, +3.6 °C. The vertical dotted line represents the separation between the rear edge (<35°N) and the expanded range. The solid line represents the significant relationship between each estimate of bolting loss and latitude, with the 95% CI indicated in gray shading. Test statistics are reported in Table S15.

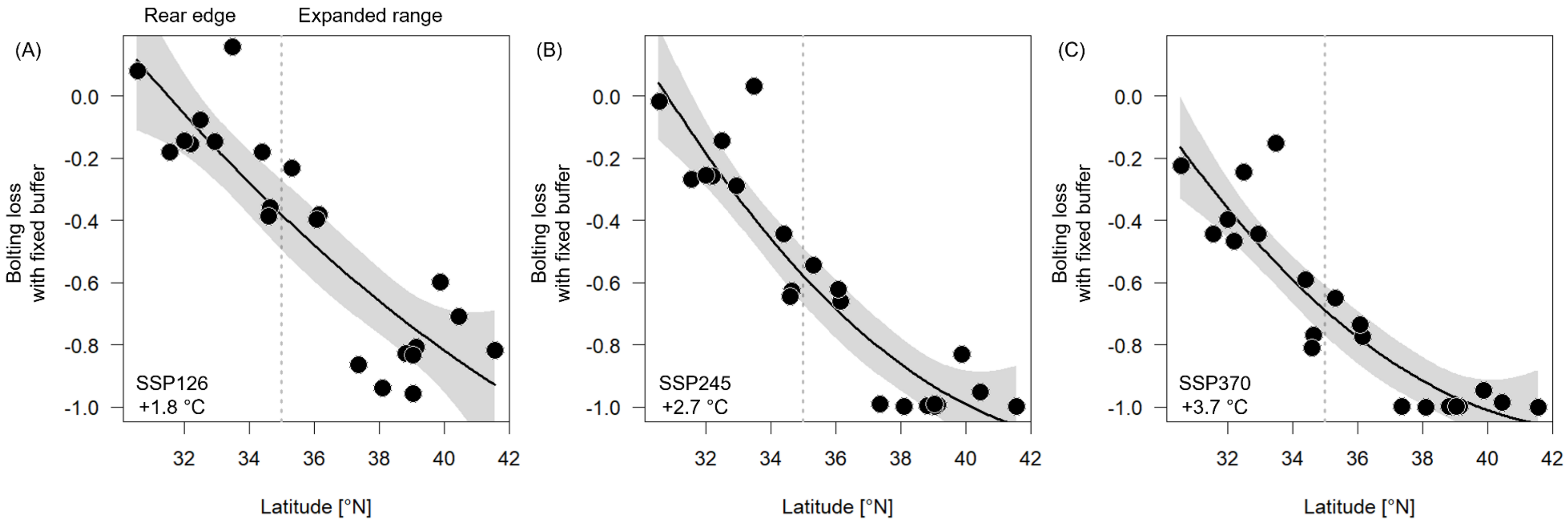

**Figure S9: Bolting loss with fixed buffer under future climates (2081–2100).** Bolting loss estimated for three Shared Socioeconomic Pathways for each population (dots), with buffering to vernalization requirement held constant across the range. **(A)** SSP126, +1.8 °C, **(B)** SSP245, +2.7 °C, **(C)** SSP370, +3.6 °C. The vertical dotted line represents the separation between the rear edge (<35°N) and the expanded range. The solid line represents the significant relationship between each estimate of bolting loss and latitude, with the 95% CI indicated in gray shading. Test statistics are reported in Table S16.

**References**

Fick, S. E., & Hijmans, R. J. (2017). WorldClim 2: New 1-km spatial resolution climate surfaces for global land areas. International Journal of Climatology, 37, 4302–4315.

Perrier, A., Turner, M. C., & Galloway, L. F. (2025). Shifts in vernalization and phenology at the rear edge hold insight into the adaptation of temperate plants to future milder winters. New Phytologist, 246, 1377–1389.

Perrier A., Keenan O. J., Busch J. W., & Galloway L. F. (2026). The legacy of past climate warming: strong local adaptation in rear-edge populations. Evolution letters, 10, 343–354.
